# A phylogenomic perspective on gene tree conflict and character evolution in Caprifoliaceae using target enrichment data, with Zabelioideae recognized as a new subfamily

**DOI:** 10.1101/2020.10.29.359950

**Authors:** Hong-Xin Wang, Diego F. Morales-Briones, Michael J. Moore, Jun Wen, Hua-Feng Wang

**Affiliations:** Key Laboratory of Tropical Biological Resources of Ministry of Education, College of Tropical Crops, Hainan University, Haikou 570228, China; Department of Plant and Microbial Biology, College of Biological Sciences, University of Minnesota, 140 Gortner Laboratory, 1479 Gortner Avenue, Saint Paul, MN 55108, USA; Department of Biology, Oberlin College, Oberlin, OH44074, USA; Department of Botany, National Museum of Natural History, MRC-166, Smithsonian Institution, PO Box 37012, Washington, DC 20013-7012, USA

**Author notes:** Authors for correspondence: Hua-Feng Wang. Authors contributed equally to this study (Hong-Xin Wang, Diego F. Morales-Briones).

**Keywords:** Caprifoliaceae, Hybridization, Introgression, Phylogenetic networks, *Zabelia*, Zabelioideae.

## Abstract

The use of diverse datasets in phylogenetic studies aiming for understanding evolutionary histories of species can yield conflicting inference. Phylogenetic conflicts observed in animal and plant systems have often been explained by hybridization, incomplete lineage sorting (ILS), or horizontal gene transfer. Here, we employed target enrichment data, species tree and species network approaches to infer the backbone phylogeny of the family Caprifoliaceae, while distinguishing among sources of incongruence. We used 713 nuclear loci and 46 complete plastome sequence data from 43 samples representing 38 species from all major clades to reconstruct the phylogeny of the family using concatenation and coalescence approaches. We found significant nuclear gene tree conflict as well as cytonuclear discordance. Additionally, coalescent simulations and phylogenetic species network analyses suggested putative ancient hybridization among subfamilies of Caprifoliaceae, which seems to be the main source of phylogenetic discordance. Ancestral state reconstruction of six morphological characters revealed some homoplasy for each character examined. By dating the branching events, we inferred the origin of Caprifoliaceae at approximately 66.65 Ma in the late Cretaceous. By integrating evidence from molecular phylogeny, divergence times, and morphology, we herein recognize Zabelioideae as a new subfamily in Caprifoliaceae. This work shows the necessity of using a combination of multiple approaches to identify the sources of gene tree discordance. Our study also highlights the importance of using data from both nuclear and chloroplast genomes to reconstruct deep and shallow phylogenies of plants.

## 1 Introduction

Gene tree discordance is a ubiquitous feature of phylogenomic data sets (Galtier and Daubin, 2008; Degnan and Rosenberg, 2009; Szöllősi et al., 2015; Sun et al., 2015; Lin et al., 2019). Many studies have shown that incomplete lineage sorting (ILS), hybridization, and other processes such as horizontal gene transfer, gene duplication, or recombination, may be contributing to discordance among gene trees (Degnan and Rosenberg, 2009; Linder and Naciri, 2015). Among these potential sources of discordance, hybridization has been especially important in plant systematics research (e.g., Morales-Briones et al., 2018; Lee-Yaw et al., 2019; Stull et al., 2020; Morales-Briones et al., 2021). Hybridization may be expected to be prevalent in rapidly radiating groups, which is increasingly recognized as a major force in evolutionary biology, in many cases leading to new species and lineages (Mallet, 2007; Abbott et al., 2010; Yakimowski and Rieseberg, 2014; Konowalik et al., 2015). ILS is one of the prime sources of gene tree discordance, which has attracted increasing attention in the past decades as phylogenetic reconstruction methods allowed its modeling (Edwards 2009; Liu et al., 2015). Despite that, distinguishing ILS from hybridization is still challenging (Linder and Naciri, 2015). More recently, methods to estimate phylogenetic networks that account simultaneously for ILS and hybridization have been developed (Solís-Lemus and Ané, 2016; Wen et al., 2018). At the same time, empirical studies using phylogenetic networks to identify the sources gene tree discordance are increasing (e.g., Morales-Briones et al., 2018, 2021; Widhelm et al., 2019; Feng et al., 2020).

Caprifoliaceae is a medium-sized family with about 960 plant species belonging to 41 extant genera that are mainly distributed in eastern Asia and eastern North America (Donoghue et al., 2001; Bell, 2004; Wang et al., 2020; Xiang et al., 2020). The family has long been the focus of phylogenetic studies of character evolution, especially regarding its tremendous diversity in reproductive structures (Backlund, 1996; Donoghue et al., 2003). Caprifoliaceae has five corolla lobes and five stamens as ancestral states, which are retained in Diervilleae C.A.Mey., *Heptacodium* Rehd., and Caprifolieae (though in some *Symphoricarpos* Duhamel and *Lonicera* L. there are four corolla lobes and four stamens). However, for other genera, the number of stamens is reduced to four or even one. Caprifoliaceae shows even greater variation in fruit types (e.g., achene in *Abelia* R. Br., berry in *Lonicera*, and capsule in *Weigela* Thunb.; Manchester & Donoghue, 1995; Donoghue et al., 2003). Some genera possess highly specialized morphological characters (e.g., the spiny leaves of *Acanthocalyx* (DC.) Tiegh.*, Morina* L. and *Dipsacus* L.) that have likely played key roles in lineage-specific adaptive radiation (Blackmore and Cannon, 1983; Caputo and Cozzolino, 1994; Donoghue et al., 2003) (Fig. 1).

**Fig. 1.**
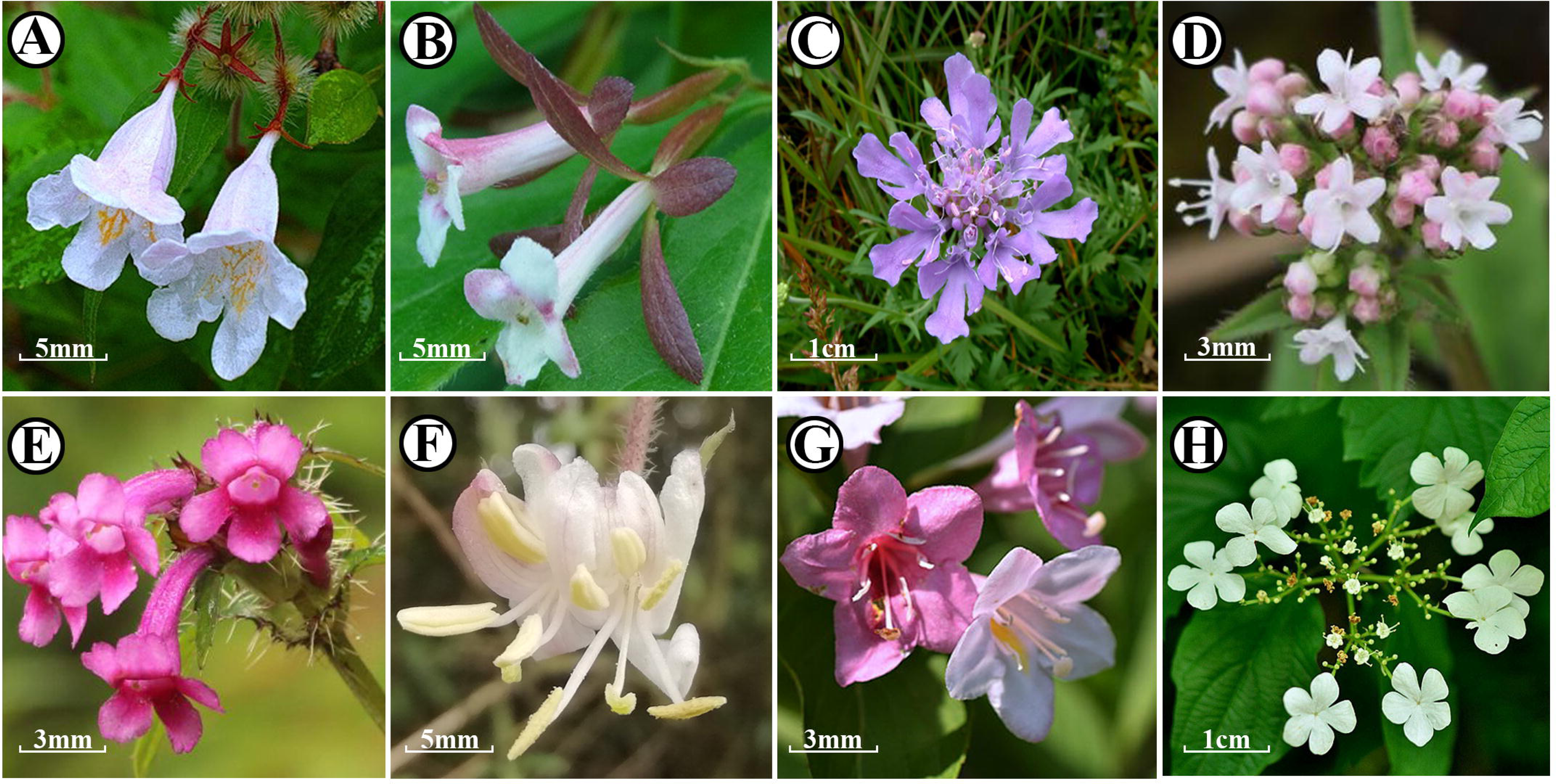
Floral diversity of Dipsacales. (A) Kolkwitzia amabilis; (B) Zabelia integrifolia; (C) Scabiosa comosa (D) Valeriana flaccidissima; (E) Acanthocalyx nepalensis subsp. delavayi; (F) Lonicera fragrantissima var. lancifolia; (G) Weigela coraeensis; (H) Viburnum opulus subsp. calvescens.

Circumscriptions of Caprifoliaceae have been controversial. Backlund & Pyck (1998) suggested that Caprifoliaceae should be defined narrowly to include only five genera, *Heptacodium* Rehder, *Leycesteria* Wall., *Lonicera*, *Symphoricarpos*, and *Triosteum* L. This narrowly circumscribed concept of the family has been also accepted by some authors (e.g., APG, 1998; Yang & Landrein, 2011; Xiang et al., 2020). By contrast, some researchers proposed to integrate Morinaceae, Dipsacaceae, Valerianaceae, and Caprifoliaceae s. str. into the Caprifoliaceae s.l. (e.g., Judd et al.,1994; Donoghue et al., 2001; Stevens, 2001 onwards; Wang et al., 2015; Wang et al., 2020). To maximize stability and ease identification based on recent phylogenetic studies (e.g., Li et al., 2019; Wang et al., 2020; Xiang et al., 2020), we prefer the Caprifoliaceae s.l. concept that includes seven major clades: Linnaeoideae, *Zabelia*, Morinoideae, Valerianoideae, Dipsacoideae, Caprifolioideae and Diervilloideae (Stevens, 2001 onwards; Wang et al., 2015; APG, 2016; Wang et al., 2020). Phylogenetic relationships within Caprifoliaceae have been studied extensively during the past two decades using plastid and nuclear DNA data (Fig. 2), but the placement of *Zabelia* (Rehder) Makino has never been resolved confidently using either morphological characters (Backlund, 1996; Donoghue et al., 2003) or molecular data (Donoghue et al., 1992; Jacobs et al., 2010; Smith et al., 2010; Landrein et al., 2012; Stevens, 2019; Wang et al., 2020; Xiang et al., 2020). Based on nuclear (ITS) and chloroplast DNA (cpDNA) data (*trnK*, *matK*, *atpB*-*rbcL*, *trnL*-*F*) of 51 taxa, Jacobs et al. (2010) found moderate support (bootstrap support [BS] = 62%) for the placement of *Zabelia* (formerly part of *Abelia*) in a clade with Morinoideae, Dipsacoideae, and Valerianoideae. Based on the same data set, Jacobs et al. (2010) raised *Abelia* sect. *Zabelia* to the genus level as *Zabelia*, and more recent studies have confirmed the distinctiveness of *Zabelia* (Landrein et al., 2012; Wang et al., 2015), often finding it sister to Morinoideae, although with low (BS ≤ 50%) to moderate (50% < BS ≤ 70%) support (Donoghue et al., 1992; Jacobs et al., 2010; Tank and Donoghue, 2010; Wang et al., 2015). Based on cpDNA data (*rbcL*, *trnL*-*K*, *matK* and *ndhF*) of 14 taxa, Landrein et al. (2012) suggested that *Zabelia* and *Diabelia* Landrein (Linnaeoideae) had similar “primitive” inflorescences of reduced simple thyrses. Landrein et al. (2012) conducted phylogenetic analyses of the Caprifoliaceae based on the structural characters of reproductive organs. In these analyses, *Zabelia* was sister to the clade of Morinoideae, and Valerianoideae + Dipsacoideae. Recently, Xiang et al. (2020) carried out analyses of complete plastomes of 32 species in this clade, demonstrating that *Heptacodium* and *Triplostegia* Wall. ex DC. are sister to Caprifoliaceae *s.s.* and Dipsacaceae, respectively, and have thus been included as members of those groups. Furthermore, *Zabelia* was found to be sister to Morinaceae in all analyses (Xiang et al., 2020). Likewise, using complete plastomes from 56 accessions representing 47 species of Caprifoliaceae, Wang et al. (2020) recovered the clade composed of Linnaeoideae, and Morinoideae + *Zabelia* as sister to Dipsacoideae + Valerianoideae) with maximum support (BS = 100%).

**Fig. 2.**
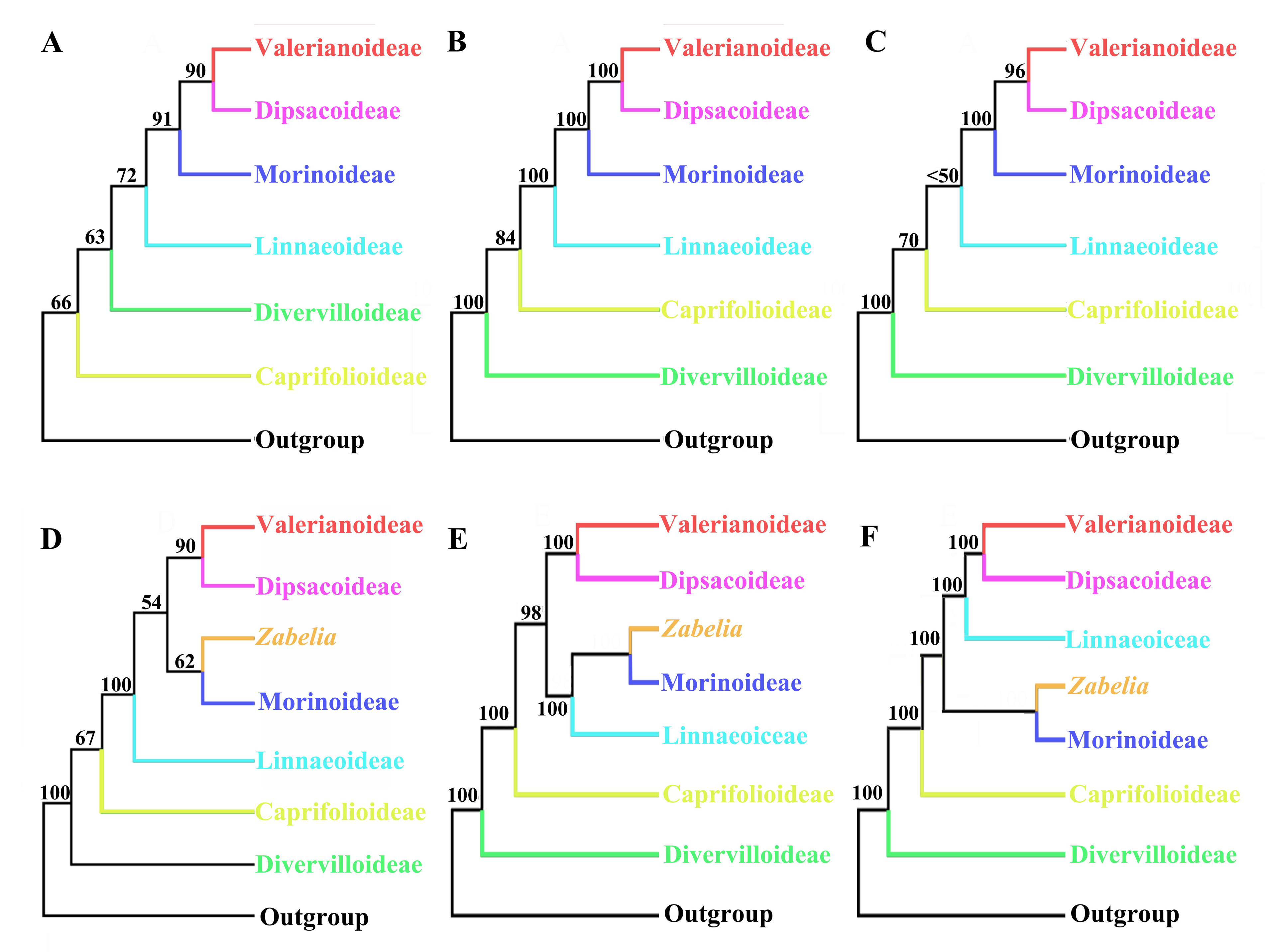
Alternative relationships for the Caprifoliaceae backbone based on previous analyses. (A) Donoghue et al. (2001); parsimony analyses based on chloroplast *rbcL* sequences and morphological characteristics; (B) Bell et al. (2001); maximum likelihood tree from the combined chloroplast DNA data; (C) Zhang et al. (2003); maximum likelihood tree based on chloroplast *trnL-F* and ndhF sequences; (D) Jacobs et al. (2010); maximum parsimony Dipsacales phylogeny based on nuclear and chloroplast sequence data; (E) Wang et al.(2020); maximum likelihood tree based on 68 complete plastomes. (F) This study; species tree based on nuclear concatenated data set.

In this study, we assembled and analyzed a custom target enrichment dataset of Caprifoliaceae to: (1) evaluate sources of gene tree discordance, in order to clarify the backbone phylogeny of Caprifoliaceae with special attention to positions of recalcitrant taxa (i.e., *Zabelia* and Morinoideae); and (2) determine the evolutionary patterns of key morphological characters of Caprifoliaceae.

## 2 Materials and methods

### 2.1 Taxon sampling

We sampled 43 individuals from 38 species of Caprifoliaceae, including representatives of all seven major clades (including *Zabelia*) of Caprifoliaceae sensu Stevens (2001 onwards) and Wang et al. (2020). Additionally, three species of Adoxaceae were included as outgroups. Most samples (38) were collected in the field, where leaf tissue was preserved in silica gel. The remaining samples were obtained from the United States National Herbarium (US) at the Smithsonian Institution (Table S1). Vouchers of newly collected samples were deposited in the herbarium of the Institute of Tropical Agriculture and Forestry (HUTB), Hainan University, Haikou, China. Complete voucher information is listed in Supporting Information Table S1.

**Table 1.**
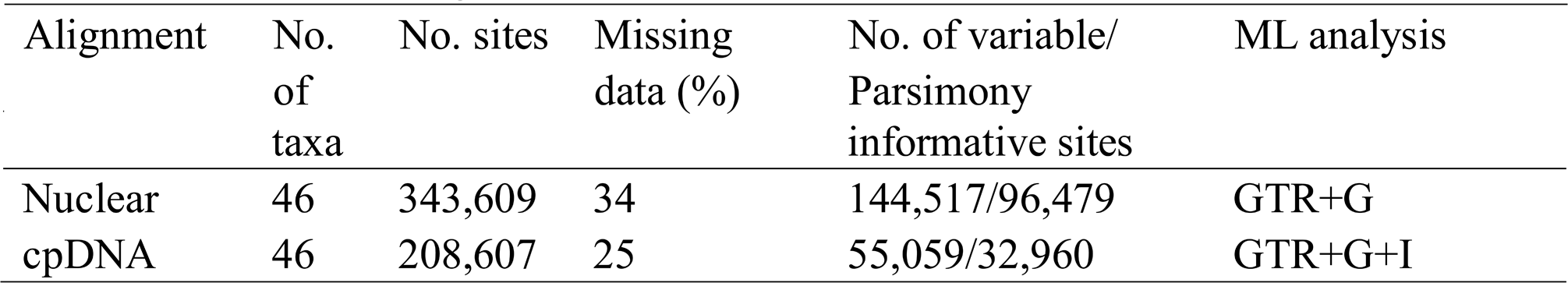
Dataset statistics, including the number of taxa, number of characters, number of PI characters, missing data.

### 2.2 DNA extraction, target enrichment, and sequencing

We extracted total genomic DNA from silica gel-dried tissue or herbarium tissue using the CTAB method of Doyle and Doyle (1987). We checked the quantity of each extraction with a Qubit 2.0 Fluorometer (Thermo Fisher Scientific, Waltham, MA, USA) and sonicated 400□ng of DNA using a Covaris S2 (Covaris, Woburn, MA) to produce fragments ∼150-350□bp in length for library preparations. To ensure that genomic DNA was sheared at approximately the selected fragment size, we evaluated all samples on a 1.2% (w/v) agarose gel.

We identified putative single copy nuclear (SCN) genes with MarkerMiner v.1.2 (Chamala et al., 2015) with default settings, using the transcriptomes of *Dipsacus asper*, *Lonicera japonica*, *Sambucus canadensis*, *Valeriana officinalis*, and *Viburnum odoratissimum* from 1KP (Matasci et al., 2014), and the genome of *Arabidopsis thaliana* (L.) Heynh. (Gan et al., 2011) as a reference. SCN genes identified with MarkerMiner were further filtered using GoldFinder (Vargas et al., 2019), requiring loci with at least 400 bp and a coverage of at least three species. This resulted in 428 SCN genes for phylogenetic analyses. A custom set of 80 bp biotinylated RNA baits (MYbaits) based on exon sequences were manufactured by Arbor Biosciences (Ann Arbor, MI, USA), with a 2× tiling density. The bait sequences are available as a supplemental file (Appendix 1).

Library preparation was done with the NEBNext Ultra II DNA Library Prep Kit for Illumina (New England Biolabs, MA, USA) following the manufacturer’s protocol. Library concentrations were quantified using a Qubit 2.0, with a dsDNA HS Assay Kit (Thermo Fisher Scientific). Fragment size distribution was determined with a High Sensitivity D1000 ScreenTape run on the Agilent 2200 TapeStation system (Agilent Technologies, Inc., Santa Clara, California, United States). Solution-based hybridization and enrichment with MYbaits followed Weitemier et al. (2014). Libraries pools were and sequenced by Novogene Corporation (Sacramento, California, U.S.A.) on one lane using the Illumina HiSeq 4000 sequencing platform (Illumina Inc, San Diego, California, U.S.A.) producing 150 □bp paired-end reads.

Given the low recovery of plastome reads from target enrichment libraries, we used a genome skimming approach to ensure recovery of full plastomes. Following Wang et al. (2020) with minor modifications, we built separate libraries for total genomic DNA. These libraries were sequenced using the BGISEQ-500 platform at BGI Shenzhen (China) with 100 bp paired-end reads.

### 2.3 Read processing and assembly

Sequencing adapters and low-quality bases were removed with Trimmomatic v0.36 (ILLUMINACLIP: TruSeq_ADAPTER: 2:30:10 SLIDINGWINDOW: 4:5 LEADING: 5 TRAILING: 5 MINLEN: 25; Bolger et al., 2014). Assembly of nuclear loci was carried out with HybPiper v.1.3.1 (Johnson et al., 2016), on an exon basis to avoid chimeric sequences in multi-exon genes that may be produced by potential paralogy (Morales-Briones et al., 2018). Only exons with a reference length of ≥ 150 bp were assembled (1220 exons from 442 genes). Paralog detection was carried out for all exons with the ‘paralog_investigator’ option of HybPiper. All assembled loci (with and without paralogs detected) were processed following Morales-Briones et al. (2020) to obtain ‘monophyletic outgroup’ (MO) orthologs (Yang & Smith, 2014).

Plastome assembly followed Wang et al. (2020). Briefly, raw reads were filtered with SOAPfilter_v2.2 (BGI-Shenzhen, China) and ddapter sequences and low-quality reads were removed. Plastome assembly was carryout using MITObim v1.8 (Hahn et al. 2013) following Wang et al. (2020).

### 2.4 Phylogenetic analyses

We used concatenation and coalescent-based methods to reconstruct the phylogeny of Caprifoliaceae. We performed phylogenetic analyses on the nuclear and plastid data sets separately. Individual nuclear exons were aligned with MAFFT v7.407 (Katoh & Standley, 2013) and aligned columns with more than 90% missing data were removed using Phyutility (Smith & Dunn, 2008). A maximum likelihood (ML) tree was estimated from the concatenated matrix, partitioning by gene, using RAxML v8.2.12 (Stamatakis, 2014) and the GTRGAMMA model for each partition. Clade support was assessed with 200 rapid bootstrap replicates (BS). We also estimated a species tree with ASTRAL-III v5.7.1 (Zhang et al., 2018) from individual ML gene trees inferred using RAxML with a GTRGAMMA model. Local posterior probabilities (LPP; Sayyari & Mirarab, 2016) were used to assess clade support.

Gene tree discordance was evaluated using two approaches. First, we mapped the individual nuclear gene trees onto the species tree and calculated the internode certainty all (ICA; Salichos et al., 2014) and number of conflicting and concordant bipartitions on each node of the species trees using Phyparts (Smith et al., 2015). Then we used Quartet Sampling (QS; Pease et al., 2018) to distinguish strong conflict from weakly supported branches in the nuclear tree. We carried out QS with 1000 replicates.

The plastomes sequences were aligned with MAFFT. A ML tree was estimated with RAxML using the GTR + I + G model and 1000 bootstrap replicates for clade support. Additionally, we used QS with 1000 replicates to evaluate branch support.

### 2.5 Assessment of hybridization

To test whether ILS alone could explain cytonuclear discordance, we used coalescent simulations similar to Folk et al. (2017) and García et al. (2017). We simulated 10,000 gene trees under the coalescent with DENDROPY v.4.1.0 (Sukumaran & Holder, 2010) using the ASTRAL species trees as a guide tree with branch lengths scaled by four to account for organellar inheritance. We summarized the simulated gene trees on the cpDNA tree. Under a scenario of ILS alone, any relationships in the empirical chloroplast tree should be present in the simulated trees and have a high frequency; under a hybridization scenario, relationships unique to the cpDNA tree should be at low (or zero) frequency (García et al., 2017).

### 2.6 Species network analysis

We inferred species networks using a maximum pseudo-likelihood approach (Yu et al., 2012). Due to computational restrictions and given our main focus on potential reticulation among major clades of Caprifoliaceae (i.e., along the backbone), except Caprifolioideae and Diervilloideae, which did not show major signal of conflict with respect to the rest of Caprifoliaceae [i.e., the remaining five major groups formed a clade with maximum support (see section 3.4)]. First, we reduced our 46-taxon data set to one outgroup and 10 ingroup taxa to include Dipsacoideae, Linnaeoeideae, Moroinoideae and *Zabelia* (11-taxa data set). To disentangle nested hybridization, we created a reduced, 9-taxa data set by removing Dipsacoideae (because *Dipsacus* and *Scabiosa* were found to be involved in several inferred hybridization events) and a 7-taxa data set that excluded these two taxa as well as *Morina* and *Zabelia* (which were found to be involved in reticulation events in both the 11-taxa and 9-taxa networks). Species network searches were carried out with PHYLONET v.3.6.1 (Than et al., 2008) with the command ‘InferNetwork_MPL’ and using individual ML gene trees. Network searches were performed using only nodes in the gene trees that had BS support of at least 50%, allowing for up to five hybridization events and optimizing the branch lengths and inheritance probabilities of the returned species networks under the full likelihood. To estimate the optimal number of hybridizations and test whether the species network fit our gene trees better than a strictly bifurcating species tree, we computed the likelihood scores of concatenated RAxML, ASTRAL and plastid DNA trees, given the individual gene trees, as implemented in Yu et al. (2012), using the command ‘CalGTProb’ in PHYLONET. Finally, we performed model selection using the Akaike information criterion (Akaike, 1973), the bias-corrected Akaike information criterion (AICc; Sugiura, 1978), and the Bayesian information criterion (Schwarz, 1978). The number of parameters equals the number of branch lengths being estimated, plus the number of hybridization probabilities being estimated, and number of gene trees used to estimate the likelihood, to correct for finite sample size.

### 2.7 Divergence time estimation

Divergence times were inferred using BEAST v.2.4.0 (Bouckaert et al., 2014). There is potential ancient hybridization in Caprifoliaceae, and therefore we estimated diversification dates separately for the nuclear and chloroplast gene tree. The root age was set to 78.9 Ma (mean 78.9 Ma, normal prior distribution 76.3–82.2 Ma) following Li et al. (2019). We selected two fossils as calibration points. First, the fossil seeds of *Weigela* Thunb. from the Miocene and Pliocene in Poland (Lańcucka-Rodoniowa, 1967), and the Miocene in Denmark (Friis, 1985) were used to constrain its stem age (offset 23.0 Ma, lognormal prior distribution 23.0 – 28.4 Ma). Second, the fruit fossil *Diplodipelta* S.R.Manchester & M.J.Donoghue, from the late Eocene Florissant flora of Colorado (Manchester, 2000; Bell & Donoghue, 2005), was used as a constraint in three different positions. In each case, the *Diplodipelta* constraint was set as an offset of 36 Ma, with a lognormal prior distribution of 34.07–37.20 Ma. Wang et al. (2015) considered three placements of *Diplodipelta* because it is possible that *Diplodipelta* represents a common ancestor of *Diabelia* and *Dipelta*, and because the sepal of *Diplodipelta* is similar to *Diabelia*, while the fruit wing of *Diplodipelta* is similar to *Dipelta* (Manchester *&* Donoghue, 2005; Wang et al., 2015). Hence, following Wang et al. (2015), we tested three placements of the *Diplodipelta* constraint: we constrained the common ancestor of *Diabelia* and *Dipelta* (Analysis I), we constrained the common ancestor (crown group) of *Dipelta* (Analysis II), and we constrained the common ancestor (crown group) of *Diabelia* (Analysis III). For each of these constraint positions, we carried out divergence time estimations for the nuclear and chloroplast trees separately.

All dating analyses were performed with an uncorrelated lognormal relaxed clock (Drummond et al., 2012), GTR + G substitution model (Posada, 2008), estimated base frequencies, and a Yule process for the tree prior. The RAxML tree was used as the starting tree, and two independent MCMC analyses of 300,000,000 generations with 10% burn-in and sampling every 3000 generations were conducted to evaluate the credibility of posterior distributions of parameters. BEAST log files were analyzed with Tracer v.1.7 (Drummond et al., 2012) for convergence with the first 10% of trees removed as burn-in. Parameter convergence was assessed using an effective sample size (ESS) of 200. Log files where combined with LogCombiner and a maximum clade credibility tree with median heights was generated with TreeAnnotator v.1.8.4 (Drummond et al., 2012).

### 2.8 Analysis of character evolution

Character states were coded from the literature, particularly from Backlund (1996), Donoghue et al. (2003), Jacobs et al. (2011) and Landrein (2017). The number of stamens was scored as follows: (0), 1; (1), 2; (2), 3; (3), 4; (4), 5. Two-character states were scored for style exertion: (0), not exceeding corolla; (1), exceeding corolla. Four fruit types were scored: (0), achene; (1), capsule, (2), berry; (3), drupe. The number of carpels was scored as: (0), 2; (1), 3; (2), 4. Number of seeds was scored as: (0), 1; (1), 2; (2), 4-5; (3), 6-20; (4), 20+; Two epicalyx types were scored: (0), no; (1), yes. All the morphological characters analyzed here are presented in Supplementary Fig. S1. Ancestral character state reconstruction was performed using ML as implemented in Mesquite v.3.51 (Maddison and Maddison, 2018) with the ‘Trace character history’ option based on the topology of the chloroplast trees. To explore differences caused by differing topologies, we also reconstructed ancestral character states onto the nuclear tree. The Markov k-state one-parameter model of evolution for discrete unordered characters (Lewis, 2001) was used.

### 2.9 Data accessibility

Raw Illumina data from sequence capture is available at the Sequence Read Archive (SRA) under accession SUB7674585 (see Table S1 for individual sample SRA accession numbers). DNA alignments, phylogenetic trees and results from all analyses and datasets can be found in the Dryad data repository.

## 3 Results

### 3.1 Assembly

The number of assembled exons per species (with > 75% of the target length) ranged from 130 (*Vesalea floribunda*) to 989 (*Diabelia sanguinea*) out of 1220 single-copy exon references, with an average of 725 exons (Table S2; Fig. S2). The number of exons with paralog warnings ranged from 1 in *Vesalea floribunda* to 619 in *Diabelia sanguinea* (Table S2). After paralog pruning and removal of exons with poor coverage across samples (at least 25 ingroup taxa), we kept 707 exons from 367 different genes. The resulting concatenated matrix had an aligned length of 343,609 bp with 96,479 parsimony-informative sites, a minimum locus size of 150 bp, and a maximum locus size of 3,503 bp, with an average of 486 bp. The plastome alignment resulted in a matrix of 208,607 bp with 32,960 parsimony-informative sites (Table 1).

**Table 2.**
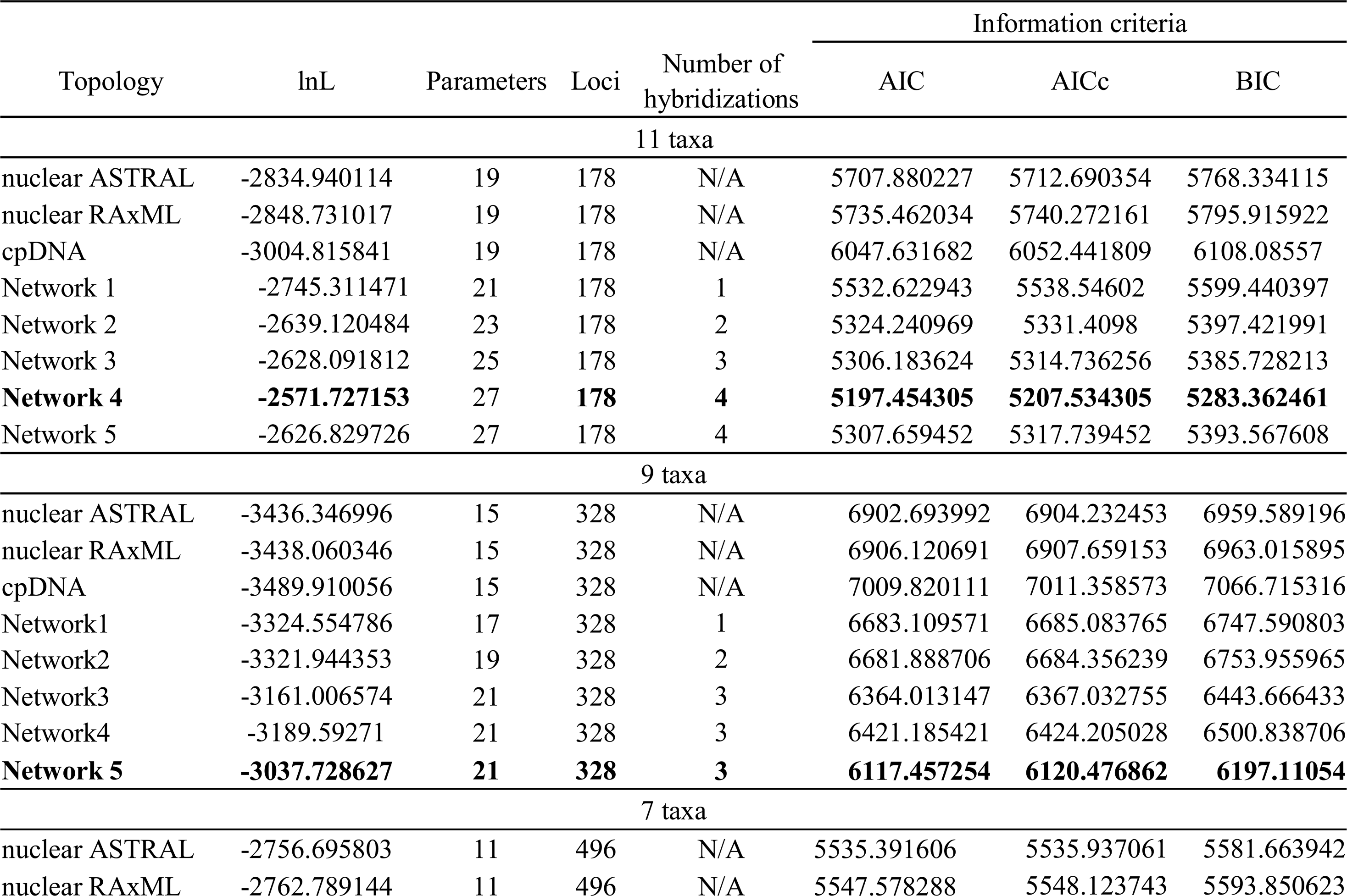
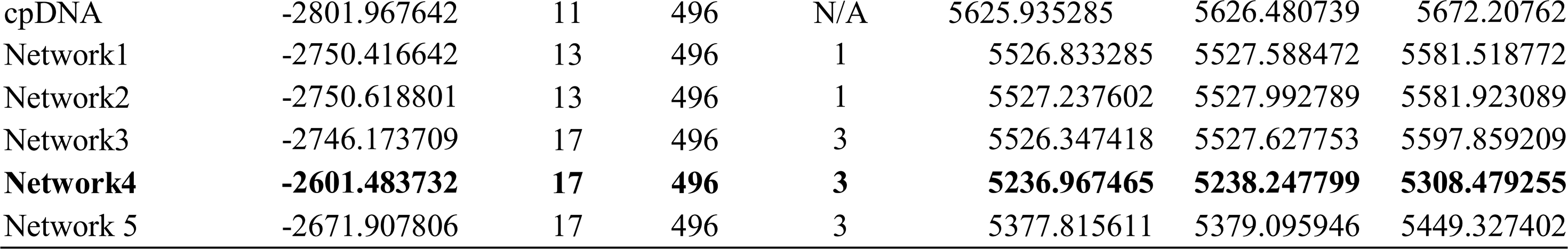
Model selection of different species networks and bifurcating trees.

### 3.2 Phylogenetic reconstruction

In analyses of both nuclear and plastid data, Diervilloideae and Caprifolioideae were successively sister to remaining Caprifoliaceae, which were resolved into five main groups: Diervilloideae, Caprifolioideae, Valerianoideae, *Zabelia* and Morinoideae. (Figs. 3–4). However, the relationships among the seven groups within Caprifoliaceae differed between nuclear and plastid analyses.

**Fig. 3.**
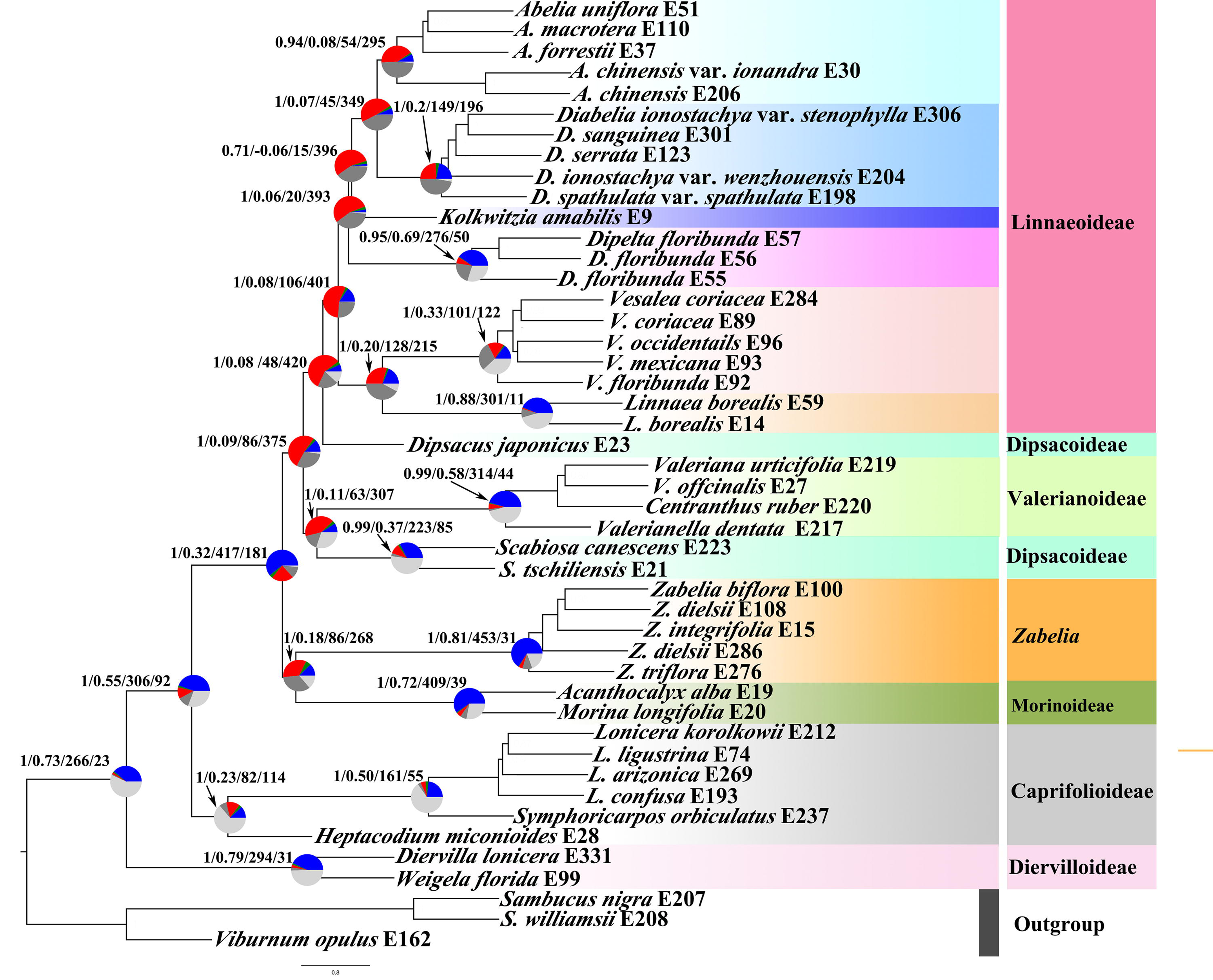
Species tree of the nuclear dataset inferred with ASTRAL-□. Local posterior probabilities and internode certainty all (ICA) scores are shown above branches for main clades. Pie charts next to the nodes present the proportion of congruent gene trees that supports that clade (blue), the proportion of discordant gene trees of the main alternative topology for that clade (green), the proportion of discordant trees for the remaining alternative topologies (red), dark gray represents the proportion of uninformative gene trees (bootstrap support < 50%), and light gray is the proportion of missing data. Numbers above branches indicate (LPP) / ICA score / number concordant gene trees / number of all discordant gene trees. Major taxonomic groups or main clades in the family as currently recognized are indicated by branch colors as a visual reference to relationships.

**Fig. 4.**
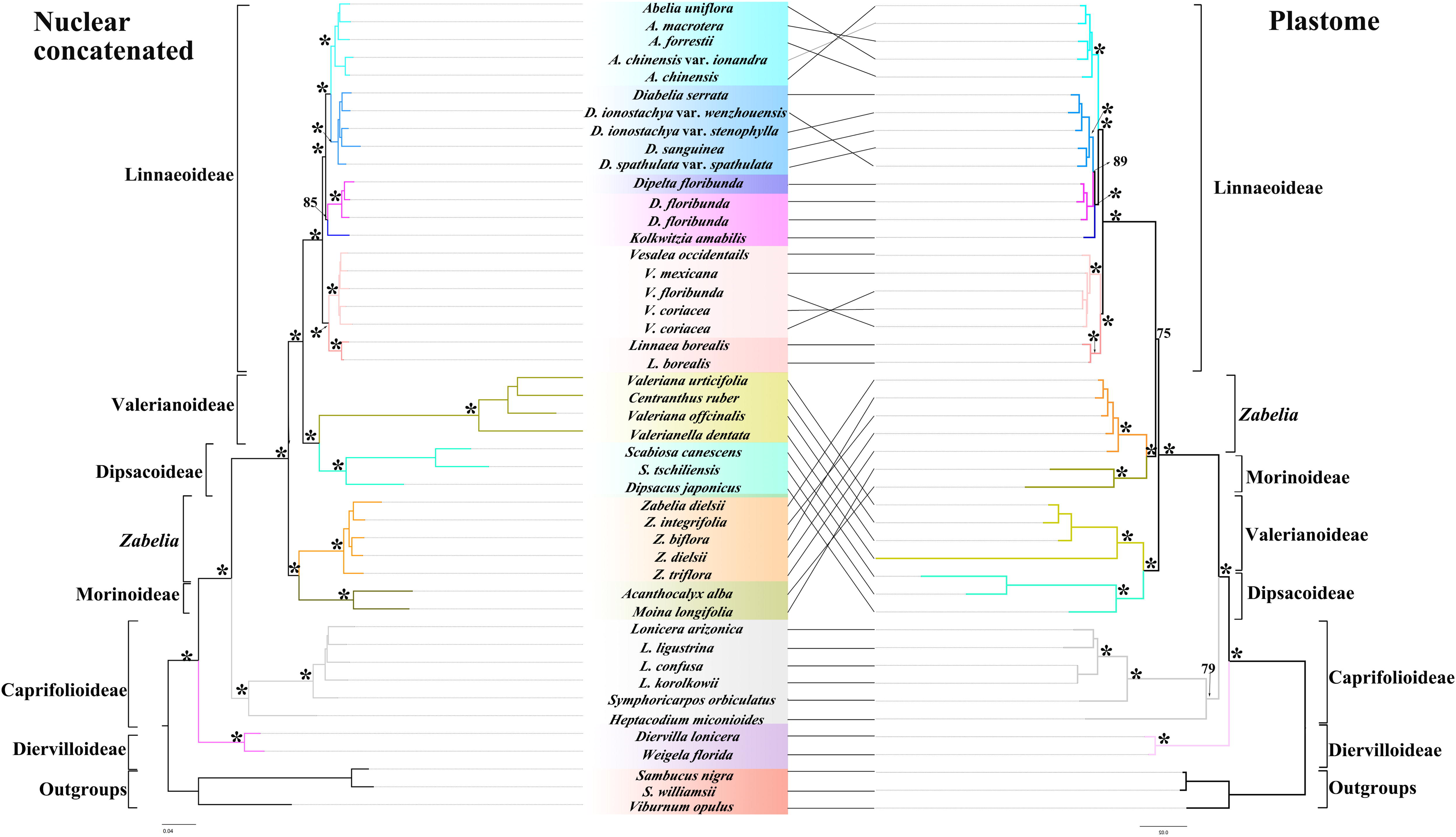
Tanglegram of the nuclear concatenated (left) and plastid (right) phylogenies. Dotted lines connect taxa between the two phylogenies. Maximum likelihood bootstrap support values are shown above branches. The asterisks indicate maximum likelihood bootstrap support of 100%. Major taxonomic groups or main clades in the family as currently recognized are indicated by branch colors as a visual reference to relationships.

#### Nuclear dataset

The ASTRAL analysis (Fig. 3) recovered maximum support (LPP = 1) for relationships within Caprifoliaceae and its seven major clades, except for the placement of *Kolkwitzia amabilis* (LPP = 0.7). Diervilloideae was resolved as sister to the rest of Caprifoliaceae, followed by Caprifolioideae as successive sister. *Zabelia* and Morinoideae formed a clade that was placed as sister to the remaining major groups. Dipsacoideae was recovered as polyphyletic, where *Scabiosa* was sister of Valerianoideae, and together with *Dipsacus japonicus* formed a grade sister to Linnaeoideae. Within Linnaeoideae, the clade of *Vesalea* M.Martens & Galeotti + *Linnaea* Gronov. ex L. was recovered as sister to a clade of all other Linnaeoideae.

The topology of the nuclear concatenated RAxML tree (Fig. 4) was mostly similar to that of the ASTRAL trees regarding major clades and their relationships. Most major clades and relationship among them had maximum support (BS = 100). The two differences were that RAxML recovered a monophyletic Dipsacoideae (*Dipascus* + *Scabiosa*) sister to Valerianoideae and *Kolkwitzia amabilis* sister of *Dipelta*.

The conflict analyses (Fig. 3, Figs. S3-S7) confirmed the monophyly of Caprifoliaceae with 266 out of 289 informative gene trees being concordant (ICA = 0.73) and having full QS support (1/–/1; i.e., all sampled quartets supported that branch). Within Caprifoliaceae, major clades and the the relationships among them had low to strong support. Diervilloideae was supported by 294 gene trees (out of 325; ICA = 0.79) and full QS support. Caprifolioideae was supported by 161 gene trees (out of 216; ICA = 0.50) and strong QS support, with signal of an alternative topology (0.87/0.077/1). The relationship of Caprifolioideae to the remaining five major clades was supported by 306 gene trees (out of 398; ICA = 0.55) and strong QS support, with signal of an alternative topology (0.82/0.043/1). The remaining five major groups formed a clade supported by 417 gene trees (out of 598; ICA = 0.32) and full QS support. Morinoideae was supported by 409 gene trees (out of 448; ICA = 0.72) and full QS support, *Zabelia* was supported by 453 gene trees (out of 484; ICA = 0.81) and also full QS support. The clade composed of *Zabelia* + Morinoideae was supported by only 86 gene trees (out of 354) and moderate QS support, with signal of a possible alternative topology (0.25/0/0.99). In the ASTRAL topology (Fig 3; Figs S3–S4), *Scabiosa* (Dipsacoideae) had 233 supporting gene trees (out of 308; ICA = 0.37) and full QS support, Valerianoideae was supported by 314 gene trees (out of 358; ICA = 0.58) with full QS support. The clade composed of Valerianoideae + *Scabiosa* was supported by only 63 gene trees (out of 370; ICA = 0.11) and moderate QS support, with signal of a possible alternative topology (0.39/0/0.99). The sister relationship of *Dipsacus japonicus* and Linnaeoideae was supported by only 48 gene trees (out of 468; ICA = 0.08) and had weak QS support, with signal of a possible alternative topology (0.073/0/1). The sister relationship of the clade Valerianoideae + *Scabiosa* and the clade *Dipsacus japonicus* + Linnaeoideae was supported only by 86 gene trees (out of 461; ICA 0.09) with moderate QS support but no signal of an alternative topology (0.17/0.73/0.97). In turn, for the RAxML topology (Figs S5 –6), a monophyletic Dipsacoideae was supported by only 80 gene trees (out of 300; ICA = 0.13) but had strong QS support with signal of an alternative topology (0.74/0/1). Linnaeoideae was supported by only 106 gene trees (out of 507; ICA = 0.08) but had strong QS support with no signal of an alternative topology (0.83/0.95/1). Within Linnaeoideae, *Linnaea* was supported by 301 gene trees (out of 312; ICA = 0.88) and full QS support, *Vesalea* was supported by 101 gene trees (out of 223; ICA = 0.33) and strong QS support but there was signal of a possible alternative topology (0.92/0/1). The clade *Linnaea +Vesalea* was supported by only 128 gene trees (out of 343; ICA = 0.20) and had strong QS support with signal of a possible alternative topology (0.92/0/1). In the case of the remaining Linnaeoideae, *Dipelta* was supported by 276 gene trees (out of 326; ICA = 0.69) and full QS support, *Diabelia* was supported by 149 gene trees (out of 345; ICA = 0.20) and had strong QS support with signal of a possible alternative topology (0.62/0/1), and *Abelia* was supported by only 54 gene trees (out of 349; ICA= 0.20) but with strong QS support and signal of a possible alternative topology (0.79/0/1). The clade formed by *Abelia + Diabelia* was supported only by 45 gene trees (out of 394; ICA = 0.07) but with strong QS support and signal of a possible alternative topology (0.67/0.2/1). In the ASTRAL analysis, the sister relationship of *Kolkwitzia amabilis* and the clade of *Abelia + Diabelia* was supported only by 15 gene trees (out of 411; ICA = −0.06) with QS counter support and clear signal for an alternative topology (−0.22/0.19/0.94). In turn, for the RAxML topology, *Kolkwitzia amabilis* was placed as sister to *Dipelta* with the support of only 39 gene trees (out of 288; ICA = 0.08) and moderate QS support, with signal for a possible alternative topology (0.22/0.29/0.95). Finally, the clade composed of *Dipelta, Kolkwitzia, Abelia,* and *Diabelia* was supported by only 20 gene trees (out of 413; ICA = 0.06) and moderate QS support with signal for a possible alternative topology (0.37/0.059/0.95).

#### Plastid dataset

Phylogenetic analysis of the cpDNA dataset also recovered the same seven major clades in Caprifoliaceae, although relationships differed between plastid and nuclear trees (Fig. 4). In the plastid tree all major clades and most relationships among them had full support (BS = 100; QS = 1/–/1; Fig. 4 and Fig. S7). The clade Valerianoideae + Dipsacoideae was recovered as sister to all remaining Caprifolicaeae. The clade composed of *Zabelia* + Morinoideae was recovered as sister to Linnaeoideae with strong BS support (75) and moderate QS support, with signal of an alternative topology (0.36/0.4/0/94). Within Linnaeoideae, *Diabelia* and *Dipelta* formed a clade with strong BS support (89) and moderate QS support, with signal of an alternative topology (0.36/0.24/0/93). *Kolkwitzia amabilis* was sister to *Diabelia* + *Dipelta* with full BS support and strong QS support, but with a signal of an alternative topology (-/94/0.33/0.99). *Abelia* was recovered with full support as the sister of the clade composed of *Diabelia*, *Dipelta* and *Kolkwitzia.* The main differences between the nuclear and cpDNA trees were the placement of *Zabelia* + Morinoideae as sister of Linnaeoideae, and the placement of *Kolkwitzia amabilis* as sister of the clade of *Diabelia* + *Dipelta*.

### 3.3 Coalescent simulations

Coalescent simulations under the organellar model did not produce gene trees that resembled the observed cpDNA tree. When the simulated gene trees were summarized on the observed chloroplast tree, most clade frequencies were near zero, as for instance *Kolkwitzia amabilis* and the clade Valerianoideae + Dipsacoideae, *Zabelia* + Morinoideae and the clade Linnaeoideae + Valerianoideae + Dipsacoideae (Fig. S8). This suggested that ILS alone cannot explain the high level of cytonuclear discordance observed in Caprifoliaceae.

### 3.4 Species network analysis

For all three data sets analyzed (11-taxa, 9-taxa, 7-taxa), any of the networks with one to five reticulations events was a better model than a strictly bifurcating tree (Table 2; Fig. S9). In the 11-taxa data set, the best network had four reticulations events, whereas three reticulation events were inferred for the best networks in the 9-taxa and 7-taxa trees (Fig. 5; Table 2). In the 11-taxa network (Fig. 5a), which included Dipsacoideae, both species of Dipsacoideae were inferred to result from hybridization events involving members of Linnaeoideae. *Dipsacus* was inferred to have contributions from three lineages, including an inferred hybridization between the lineages leading to *Linnaea* and *Vesalea*. In the 9-taxa tree (excluding Dipsacoideae but including *Morina* and *Zabelia*; Fig. 5b), both *Morina* and *Zabelia* were inferred to have genetic contributions from Linnaeoideae, while the clade of *Abelia* + *Diabelia* was inferred to have received a contribution from the lineage leading to *Kolkwitzia*. In the 7-taxa tree (including only Linnaeoideae + *Viburnum*; Fig. 5c), the clade of *Abelia* + *Diabelia* was again inferred to have resulted from a reticulation event, as were *Dipelta* and *Linnea*, although in the case of *Linnea*, there was only a small contribution (1.2%) from the lineage leading to *Vesalea*.

**Fig. 5.**
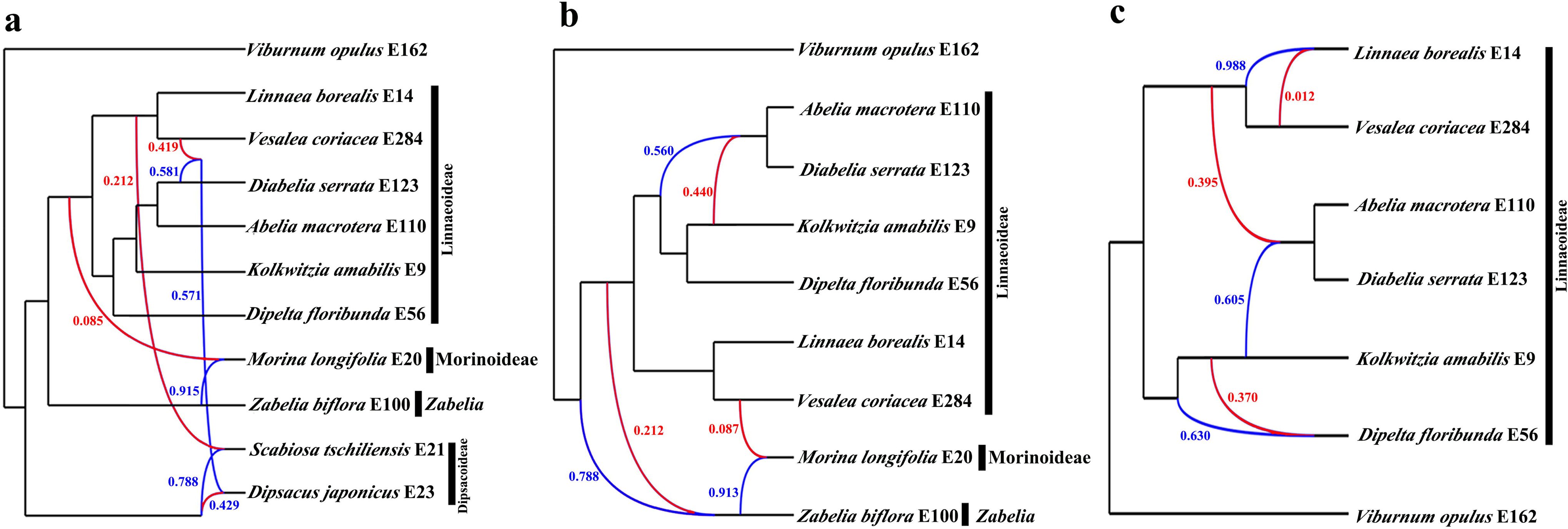
Best supported species networks inferred with Phylonet for the (a) 11-taxa, (b) 9-taxa, and (c) 7-taxa data sets. Numbers next to the inferred hybrid branches indicate inheritance probabilities. Red lines represent minor hybrid edges (edges with an inheritance contribution < 0.50)

### 3.5 Divergence time estimation

Divergence time estimates based on the nuclear data set suggested that the deepest divergences in Caprifoliaceae occurred in the early Paleocene, whereas most generic-level diversification occurred in the middle Eocene to middle Oligocene (Fig. 6). The divergence between Dipsacoideae and Valerianoideae was dated to 41.83 Ma (95% Highest Posterior Density (HPD) = 34.44–49.35 Ma) (Fig. 6). The diversification of Linnaeoideae was inferred to be at 48.33 Ma (95% HPD = 42.37–51.70 Ma) (Fig. 6). Within Linnaeoideae, both *Abelia* and *Kolkwitzia* originated almost contemporaneously in the mid-late Eocene. The onset of *Zabelia* and Morinoideae diversification occurred between 32.10 and 39.89 Ma (Fig. 6). A comparison of the time estimates for selected nodes under the six different analyses is available in Figs. S10 and S11. As would be expected, Analysis I (placing the *Diplodipelta* fossil constraint at the ancestor of *Diabelia* and *Dipelta*) resulted in younger ages for much of Linnaeaoideae, but otherwise did not significantly affect divergence dates. We found that our estimated ages were generally younger with nuclear vs. plastid data (Figs. S10 - S11). For instance, our analyses showed that Caprifoliaceae date to 66.65 Ma (95% HPD = 56.31-69.44 Ma) in analysis I based on nuclear datasets (Fig. S 10), whereas in the two other analyses, the divergence time was estimated as 69.38 Ma (95% HPD = 53.26-79.45 Ma) and 69.87 Ma (95% HPD = 64.20-76.34 Ma, Fig. S10). Based on chloroplast datasets, the divergence time of Caprifoliaceae was estimated at 76.43 Ma (95% HPD = 64.81-82.10 Ma), 78.61 Ma (95% HPD = 71.33-82.45 Ma) and 77.43 Ma (95% HPD = 58.33-81.60 Ma) in the three analyses respectively (Fig. S10).

**Fig. 6.**
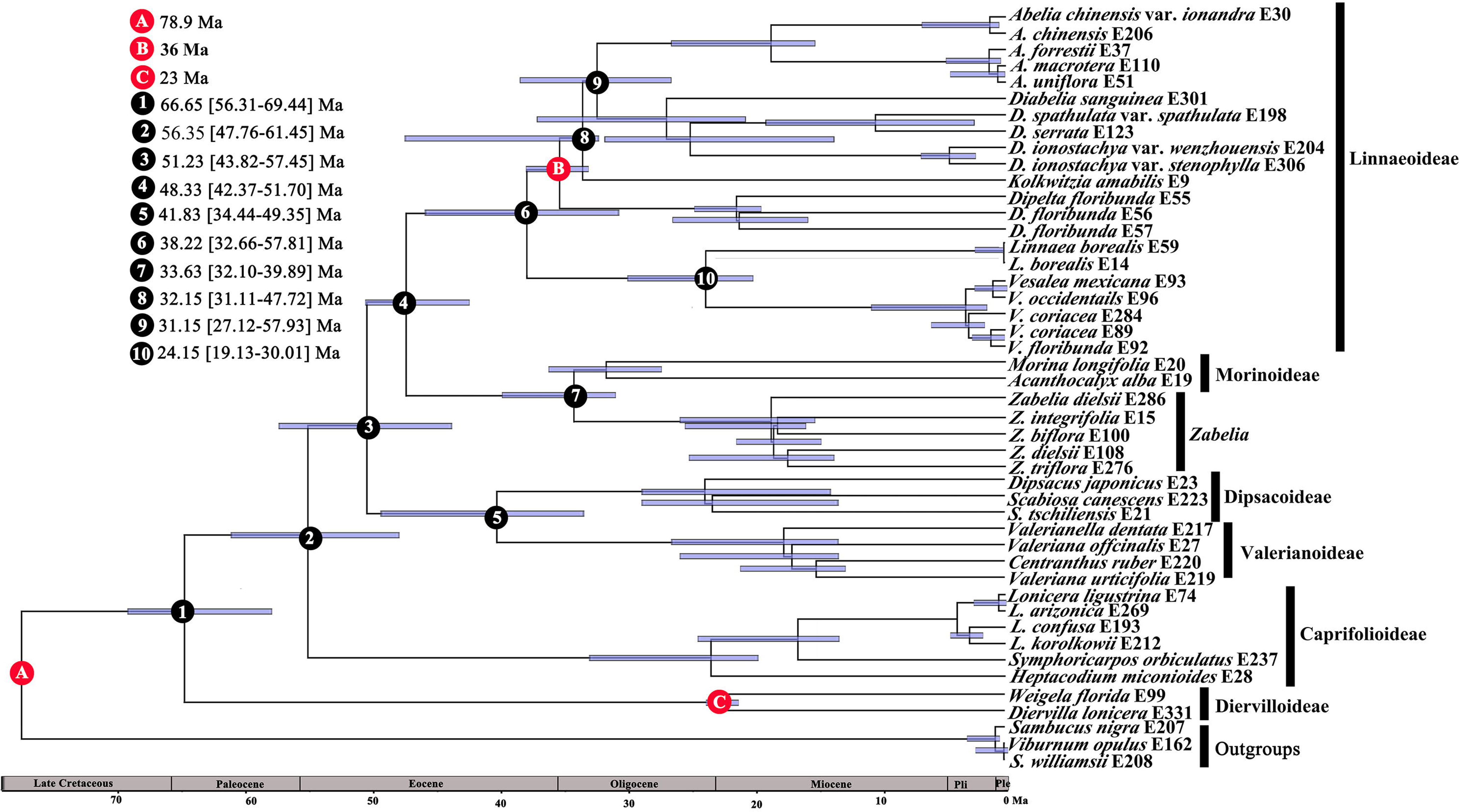
BEAST analysis of divergence times based on the nuclear data set, under Analysis I. Calibration points are indicated by A, B. and C. Numbers 1–11 represent major divergence events in Caprifoliaceae; mean divergence times and 95% highest posterior densities are provided for each nodes of interests.

### 3.6 Character evolution

The likelihood inference of character evolution using the cpDNA tree detected some homoplasy in each of the six morphological characters examined (Figs. 8 - 10), with style exertion relative to corolla showing particularly high homoplasy (Fig. 7). Stamen number exhibited little homoplasy, with an inferred shift from five stamens ancestrally to four in the bulk of Caprifoliaceae, and within Valerianoideae, it further reduced to 3 and 1 (Fig. 7). In contrast, style exertion exhibited a high level of homoplasy in the early diversification of the family (Fig. 7). Even within Linnaeoideae, the state “not exceeding corolla” was inferred to have originated twice, in *Vesalea* and in the *Diabelia*-*Dipelta*-*Kolwitzia*-*Abelia* clade. The ancestral fruit type for Caprifoliaceae was uncertain due to the diversity in fruit types among the major early-diverging clades, but was most likely an achene (Fig. 8). Carpel number evolution was similarly complex (Fig. 8), with three carpels inferred to be ancestral in the family. For seed number, one seed was inferred as the ancestral character state for Caprifoliaceae, with independent shifts to two seeds in *Symphoricarpos* and *Dipelta* (Fig. 9). Independent origins of the epicalyx in *Dipsacus* and the clade of *Moina* and *Acanthocalyx* was also inferred (Fig. 9).

**Fig. 7.**
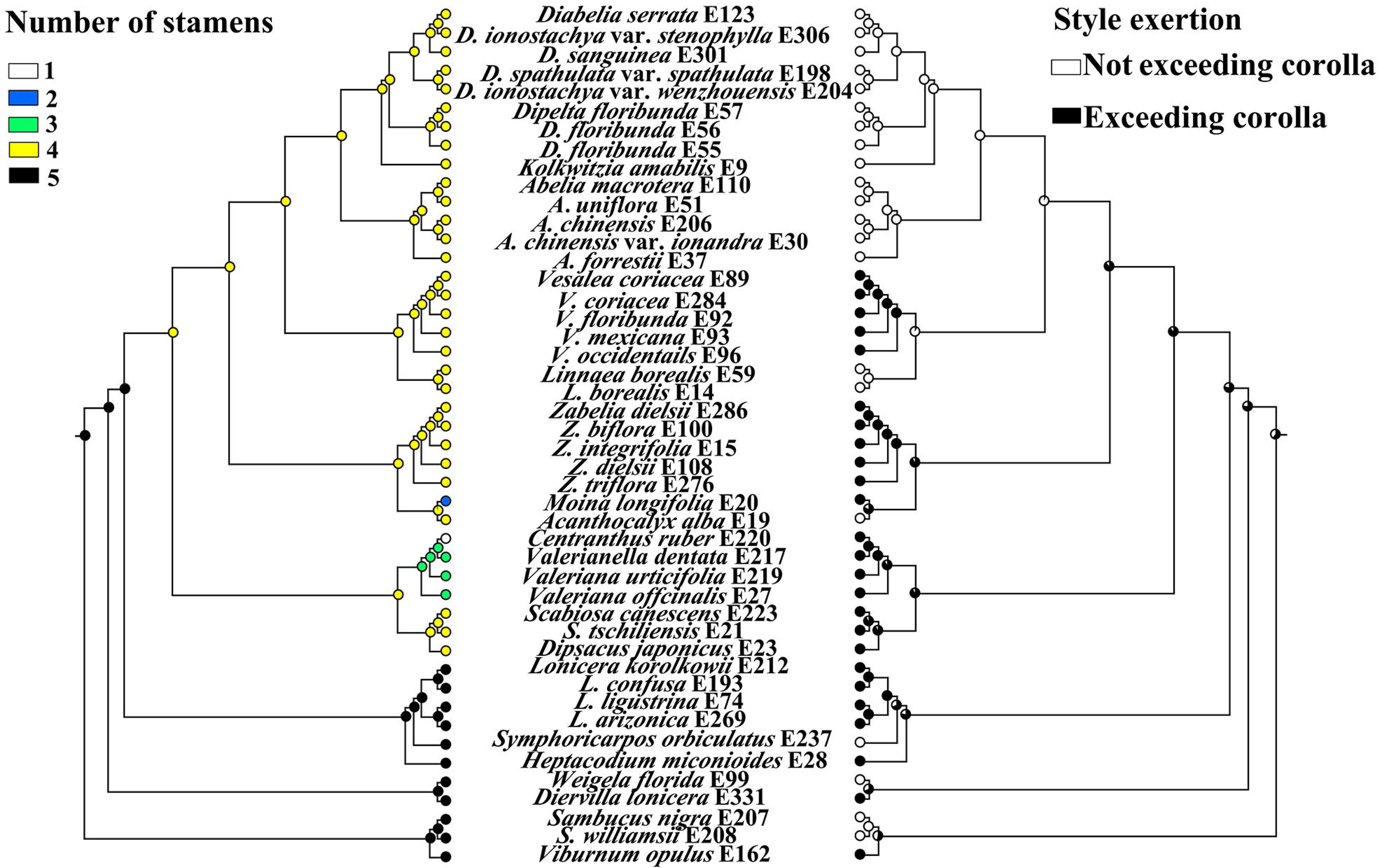
Maximum likelihood inference of character evolution in Caprifoliaceae based on the plastid matrix. Left, Number of stamens; Right, Style exertion.

**Fig. 8.**
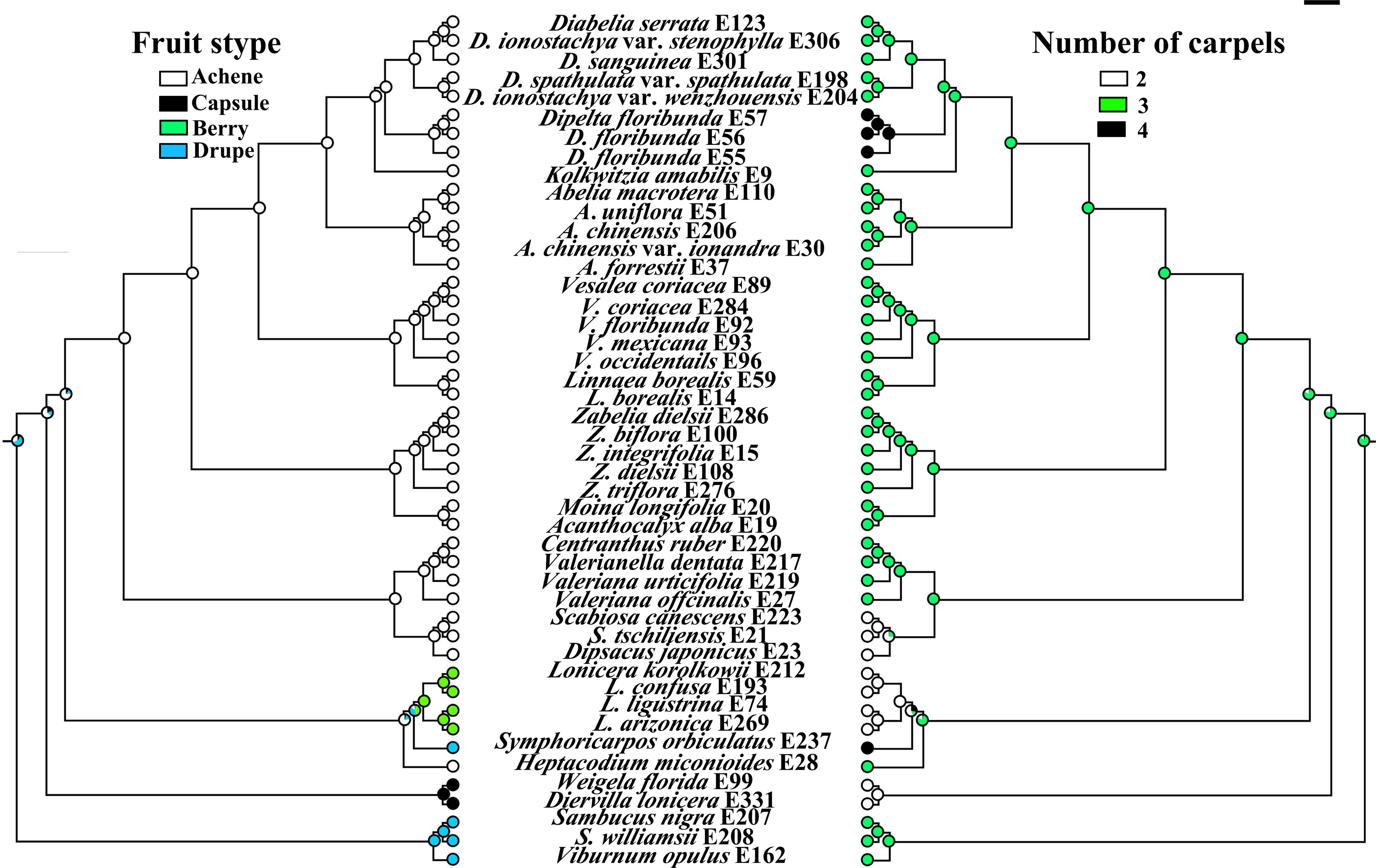
Maximum likelihood inference of character evolution in Caprifoliaceae based on the plastid matrix. Left, fruit type; Right, Number of carpels.

**Fig. 9.**
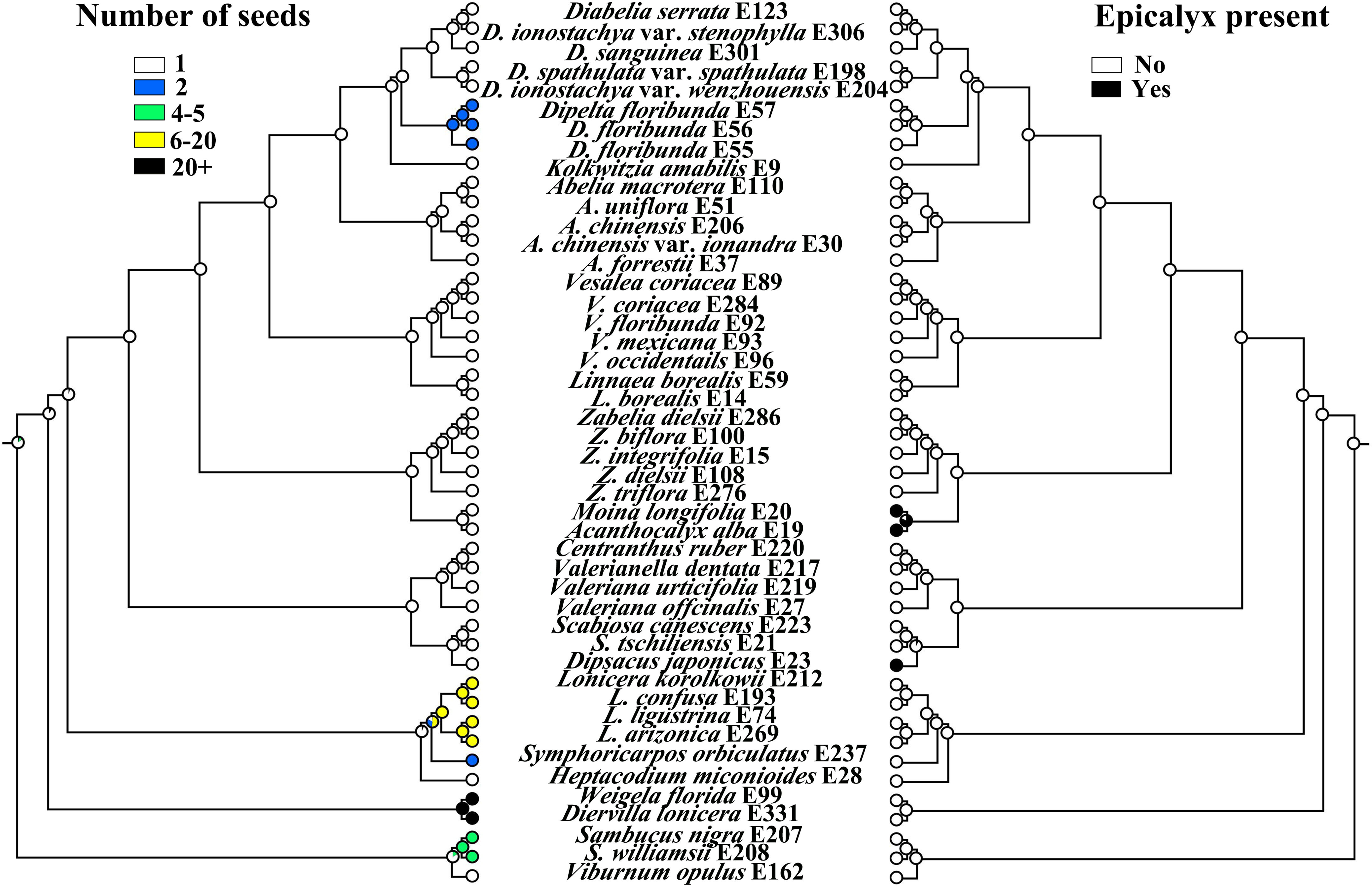
Maximum likelihood inference of character evolution in Caprifoliaceae based on the plastid matrix. Left, number of seeds; Right, epicalyx presence/absence.

A summary of character state evolution using the nuclear gene tree and character states that are relevant for the taxonomy of the group is shown in Figs. S12, S13 and S14. We found that the patterns of character evolution from cpDNA tree and nuclear gene tree were similar.

## 4 Discussion

### 4.1 Phylogenetic incongruence and putative hybridization

Although both our nuclear and plastid phylogenies supported the same seven major clades of Caprifoliaceae, the relationships among these clades are incongruent between data sets (Figs. 5 and 6). For instance, in the nuclear ASTRAL tree, Linnaeoideae is recovered as sister to Dipsacoideae (except for *Dipsacus japonicus*) + Valerianoideae (Fig. 3), while in the plastid tree Linnaeoideae is sister to *Zabelia* + Morinoideae (Fig. 4). In contrast, in the nuclear RAxML concatenated tree (Fig. 3), Linnaeoideae is recovered as sister to Dipsacoideae +Valerianoideae. Some of these points of conflict pertain to areas of Caprifoliaceae phylogeny that have long been problematic—for example, the relationships between *Zabelia* and other subfamilies. The inclusion of extensive nuclear genome sampling for Caprifoliaceae in this study is important because plastome-only phylogenies may not fully capture evolutionary processes such as ILS or organellar capture via hybridization. Three main processes will lead to gene tree heterogeneity and cytonuclear discordance: gene duplication/extinction, horizontal gene transfer/hybridization, and ILS. Currently, there are many methods to detect gene discordance (e.g., Smith et al., 2015; Pease et al., 2018), but sources of such discordance remain hard to disentangle, especially when multiple processes co-occur (e.g., Morales-Briones et al. 2021).

*Zabelia* was long thought to be closely related to *Abelia* (Hara, 1983; Tang & Lu, 2005). However, based on molecular datasets, Tank and Donoghue (2010) and Jacobs et al. (2011) found that *Zabelia* was sister to Morinaceae (=Morinoideae in this study). Using six molecular loci and inflorescence morphology, Landrein et al. (2012) concluded that the position of *Zabelia* remained unclear. The molecular investigation of Xiang et al. (2020) found that the sister relationships between *Zabelia* + Morinaceae and Linnaceae + Valerianaceae + Dipsacaceae were not highly supported. Such phylogenetic incongruence provides the opportunity to test causal hypotheses of cytonuclear discordance, e.g., ILS or hybridization. Further, in our analyses (Fig. 4), widespread cytonuclear discordance exists across Caprifoliaceae, especially at genus levels, with a high level of conflict within genera. Regarding deep Caprifoliaceae relationships, the results from the nuclear analyses (Figs. 4 and 5) showed multiple instances (at least two) of well-supported conflict with the results from the plastome (Fig. 4), and the plastid results were largely consistent with previous plastid and large-scale analyses of Caprifoliaceae (Wang et al., 2020).

It is worth mentioning that Dipsacoideae was not recovered as monophyletic only in the ASTRAL species tree (Fig. 3), in which *Dipsacus japonicus* had a sister relationship with Linnaeoideae. Still, nodes with the strong LPP support also had low ICA and QS support values, which suggests that ILS and/or unidentified hybrid lineages continue to obscure our understanding of relationships in Dipsacoideae.

Previous studies reported that hybridization has shaped the evolutionary history of Caprifoliaceae (e.g., *Heptacodium miconioides*) (Zhang et al., 2003; Landrein et al., 2002). The conflict analyses of the nuclear dataset revealed strong signals of gene tree discordance among the seven major clades of Caprifoliaceae. The coalescent simulations also suggested that the observed cytonuclear discordance cannot be explained by ILS alone, which along with the phylogenetic network analyses point to several potential reticulation events along the backbone of Caprifoliaceae (Fig. 5). *Morina* and *Zabelia* are frequently involved in inferred reticulations, and the two Dipsacoideae are involved in 3 of the four events in the 11-taxa tree. The clade of *Abelia* + *Diabelia* are also involved in inferred reticulation events in the 9-taxa and 7-taxa networks. If these inferred events correspond to actual past instances of gene flow (which can only be confirmed by more detailed genomic analyses), it would help to explain the high amount of phylogenetic conflict observed in our analyses. There is some potential morphological support for ancient hybridization. For example, the leaves of *Morina* have stiff spines, while the leaves of *Zabelia* have no spines. In part because of this, Wang et al. (2015) suggested that *Zabelia* may be of allopolyploid origin.

### 4.2 Temporal divergence of Caprifoliaceae

Our estimated ages using nuclear and chloroplast trees are generally younger than those of Wang et al. (2015) and Wang et al. (2020) based on two reliable fossils (Li et al., 2019). We found that the diversification and global spread of the subfamilies of Caprifoliaceae occurred during the late Cretaceous, Paleocene and Eocene (Figs. 6, S10, S11), similar to the results of Beaulieu et al. (2013). The divergence times of Caprifoliaceae have been estimated to be around the Cretaceous–Paleogene (K-Pg) boundary (Figs. 6, S10). Our results are congruent with the phenomena reported in several other plant groups such as Amaranthaceae *s.l.* (Morales-Briones et al. 2021) and legumes (Koenen et al., 2020), and in lichenized fungi such as Lobariaceae (Ascomycota) (Widhelm et al., 2019). It is generally accepted that because of the mass extinctions that occurred around the K-Pg boundary, new habitats became available and diverse organisms experienced rapid diversifications (Schulte et al., 2010). As a result of later tectonic movements and climate fluctuations from the Paleocene to the Eocene, major Caprifoliaceae lineages subsequently underwent rapid diversifications.

The divergence times among the major lineages of the Caprifoliaceae were dated to the Oligocene and Eocene, and within-genus diversification was dated to the Miocene and Pliocene (Figs. 6, S10, S11). Our results may be explained by the hypothesis that members of the Caprifoliaceae are well adapted to relatively cool environments (Friis, 1985; Manchester and Donoghue, 1995; Manchester, 2000), and an increase in the earth’s temperature in the late Paleocene and early Eocene may have forced them to move to higher elevations or latitudes. As plants moved to higher elevations, their distribution was likely to be fragmented, resulting in isolation between populations. We have some evidence to support this hypothesis: (1) This family is mainly distributed in north temperate zones, and some genera even reach areas near the Arctic Circle (such as *Linnaea*); (2) there are numerous species (such as *Valeriana officinalis*, *Lonicera rupicola*, and *L. spinos*a) with island-like distributional patterns at relatively high elevations. As global climates cooled beginning in the late Eocene and especially the Oligocene and Miocene, cold-adapted survivors of warmer climates may have flourished and shifted into new geographic areas, especially mountainous areas, but may have struggled in northern regions during the Pliocene and Pleistocene glacial cycles (Moore & Donoghue, 2007). These global climatic events (e.g., ancient orogeny and monsoon-driven events) that might have driven diversification in Caprifoliaceae have also been reported in other taxa (Lu et al., 2018; Ding et al., 2020). For example, some genera or taxa with tiny, narrow or needle-like leaves (e.g., *Linnaea*, *Lonicera myrtillus*) may have benefited from the global cooling and drying of the Miocene and Pliocene by expanding their ranges, while other lineages more adapted to the wetter, warmer parts of the world (*Abelia, Diabelia,* and *Dipelta*) may not have contracted during the same time period.

### 4.3 Evolution of morphological characters

Character state reconstruction was conducted using ML (Figs. 8-10) because of the potential hemiplasy and xenoplasy produced by the discordance and hybridization detected in the nuclear backbone (Avies & Robinson, 2008; Robinson et al., 2008; Copetti et al., 2017; Wang et al., 2020). A consequence of this discordance may be elevated levels of apparent homoplasy in the species tree (Copetti et al., 2017; Hahn & Nakhleh, 2017).

Stamen number, fruit type, style exertion, number of carpels, number of seeds and epicalyx presence have been traditionally used for generic recognition within Caprifoliaceae (Backlund, 1996; Donoghue et al., 2003; Yang & Landrein, 2011; Landrein et al., 2020). Discordance among morphological traits might plausibly arise due to either variable convergent selection pressures or other phenomena such as hemiplasy. The evidence indicates that the probability of hemiplasy is high for four morphological characters in Caprifoliaceae: the branch lengths leading to lineages with derived character states are uniformly short with high levels of gene tree discordance. It is possible that gene flow has contributed to these patterns. For example, the ancestral stamen number states (i.e., 2 and 4) found in *Morina longifolia* and *Acanthocalyx alba* within the Morinoideae clade could be due to introgressed alleles, as we identified putative introgression between those lineages (Fig. 5). Morphological and anatomical studies showed that the earliest Caprifoliaceae had monosymmetric flowers (probably weakly so at first) with larger calyx lobes, tubular corollas, elongate styles, and capitate stigmas (Donoghue et al., 2003). Within Caprifoliaceae, the main change in stamen number is a reduction from five to four stamens. Subsequently, there was a reduction to two stamens within Morinoideae and to three, two, and one within Valerianoideae (Figs. 8 and S11). These variations may be related to an underlying change in floral symmetry (Donoghue et al., 2013), which may relate to carpel abortion or to differences in the arrangement of flowers at the level of the inflorescence.

Our results suggest that multiple independent evolutionary events of the carpel evolution in Caprifoliaceae have occurred (Figs. 9 and S12). In Caprifoliaceae, the abortion of two of the three carpels and the development of a single ovule within the remaining fertile carpel was evidently correlated with fruit type (Wilkinson, 1949). For some subfamilies of Caprifoliaceae, carpel abortion occurs at a relatively late stage of ovary development, so many species have two empty chambers at fruit maturity (e.g., Linnaeoideae, Morinoideae, and Valerianoideae). In fact, in some species, these empty compartments have been co-opted in various ways in connection with dispersal (e.g., inflated for water dispersal in some *Valeriana*).

Caprifoliaceae shows great variation in fruit types. Fleshy, bird-dispersed fruits are limited to the Caprifolioideae (Donoghue et al., 2003). *Lonicera* has berries, though generally with just a few seeds embedded in copious pulp. Based on our analyses, it is important to note that the ancestral carpel number for Caprifoliaceae is most likely 3. There is programmed carpel abortion and the number of seeds corresponds to the number of fertile carpels (Donoghue et al., 2003). For *Symphoricarpos*, two of the four carpels abort, and there are two stones. The mesocarp in the cases is rather dry and mealy in texture. In the Caprifoliaceae, achenes with a single seed are present in *Heptacodium* and in the large Linnaeoideae clade (though in *Dipelta*, and in *Linnaea* there are two seeds at maturity). From the standpoint of fruit evolution, the linkage of *Heptacodium* within Caprifolioideae implies either the independent evolution of achenes or a transition from achenes to fleshy fruits in the line leading to Caprifolioideae. Among the achene-producing Caprifoliaceae, there are various adaptations for wind dispersal. One of the most striking of these modifications is enlargement of the calyx lobes into wings as the fruits mature (e.g. in *Abelia, Dipelta* and *Diabelia*). Especially well known is the production of a feathery pappus-like structure in species such as *Valeriana officianalis* and *Centranthus ruber* in Valerianoideae. This modification facilitates passive external transport by animals. A similar case is also found in *Kolkwitzia*.

The reconstruction of character evolution thus shows that som<u>e</u> characters that were once considered important for taxonomy within the family have been inferred to be the results of homoplasious evolution (Gould 2000; Pyck, 2001; Bell 2001, 2004; Carlson et al., 2009; Zhai et al., 2019). In analysis of character evolution, homoplasy is regarded as noise that, if not properly accommodated, jeopardizes phylogenetic reconstructions using morphological characters. At the same time, hemiplasy is one of the causes of homoplasy (Copetti et al., 2017). The phenomenon of hemiplasy is most plausible when the internodal distances in a phylogenetic tree are short (relative to effective population sizes) (Robinson et al., 2008). Furthermore, the extensive hybridization detected in the backbone of Caprifoliaceae might further contribute to hemiplasy and xenoplasy (Wang et al. 2020). This may explain why it has been difficult to reconstruct the relationships and character evolution among the major lineages and genera of the family. Eventually, more extensive sampling and developmental studies will be needed to elucidate the mechanisms underlying the morphological evolutionary patterns outlined here.

### 4.4 Recognition of Zabelioideae as a new subfamily in Caprifoliaceae

Despite the strong signals of gene tree discordance, our nuclear and plastid phylogenies strongly support seven major clades in Caprifoliaceae: Linnaeoideae, *Zabelia*, Morinoideae, Valerianoideae, Dipsacoideae, Caprifolioideae and Diervilloideae, and show *Zabelia* as the sister to the morphologically highly distinct Morinoideae (Figs. 4 - 5). Our analyses support reticulate evolution concerning the origins of both the *Zabelia* lineage as well as the Morinoideae. Based on the phylogenomic and morphological analyses, we herein propose to recognize *Zabelia* as representing a new subfamily of Caprifoliaceae.

**Zabelioideae B. Liu & S. Liu ex H.F. Wang, H.X. Wang, D.F. Morales-B, M.J. Moore & J. Wen, subfam. nov.**

**Type: *Zabelia* (Rehder) Makino.**

#### Description

Shrubs, deciduous; old branches often with six deep longitudinal grooves. Leaves opposite, entire or dentate at margin; estipulate; petioles of opposite leaf pairs dilated and connate at base, enclosing axillary buds. Inflorescence a congested thyrse of cymes; cymes 1-3-flowered. Calyx 4- or 5-lobed, persistent, spreading. Corolla 4- or 5-lobed, hypocrateriform, ± zygomorphic; corolla tube cylindrical. Stamens 4, included, didynamous. Ovary 3-locular, 2 locules with 2 series of sterile ovules and 1 locule with a single fertile ovule; stigmas green, capitate, mucilaginous. Fruit an achene crowned with persistent and slightly enlarged sepals. Basic chromosome number *x* = 9.

One genus and six species distributed in China, Japan, Korea, Afghanistan, NW India, Kyrgyzstan, Nepal, and Russian Far East.

Zabelioideae is highly distinct morphologically from its sister Morinoideae. They can be easily distinguished by their habit (with Zabelioideae as shrubs, and Morinoideae as herbs), the six distinct, longitudinal grooves on twigs and branches of Zabelioideae (the six grooves absent in Morinoideae), and the epicalyx (absent in Zabelioideae and present in Morinoideae). Zabelioideae and Morinoideae share some similarities in pollen micromorphology, as both have psilate pollen grains with an endocingulum (Verlaque, 1983; Kim et al., 2001; Jacobs et al., 2011). The two subfamilies diverged in the early-mid Eocene (Figs. 6, S7), and their long evolutionary history associated with deep hybridization events, ILS and extinctions likely have made it difficult to determine their phylogenetic placements.

## 5 Conclusions

Gene tree discordance has been commonly observed in phylogenetic studies. Moreover, species tree estimation has been shown to be inconsistent in the presence of gene flow (Solís-Lemus et al., 2016; Long & Kubatko, 2018), which suggests that both ILS and gene flow simultaneously need to be considered in constructing phylogenetic relationships. Here, our results show clear evidence of cytonuclear discordance and extensive conflict between individual gene trees and species trees in Caprifoliaceae. We also show that there has been widespread hybridization and/or introgression among the major clades of Caprifoliaceae, which can explain most the gene tree conflict and the long history of phylogenetic uncertainty in the family. Furthermore, the temporal diversification of Caprifoliaceae provides a good case to support the evolutionary radiation of a dominantly north temperate plant family in response to climatic changes from the late Cretaceous to the present. Finally, based on evidence from molecular phylogeny, divergence times, and morphological characters, we herein recognize the *Zabelia* clade as representing a new subfamily, Zabelioideae, in Caprifoliaceae. The phylogenetic framework presented here also sheds important insights into character evolution in Caprifoliaceae.

## Acknowledgements

The work was funded by National Scientific Foundation of China (31660055). We thank Gabriel Johnson for his help with the target enrichment experiment, and the United States National Herbarium for permission to sample some collections. We acknowledge the staff in the Laboratories of Analytical Biology at the National Museum of Natural History, the Smithsonian Institution for support and assistance.

## Author contributions

H.F.W. and J.W. conceived the study. H.F.W. and D.F.M-B. performed the research and analyzed the data. H.X.W., H.F.W., D.F.M-B., J.W. and M.J.M wrote and revised the manuscript.

---

**Fig. S1.**
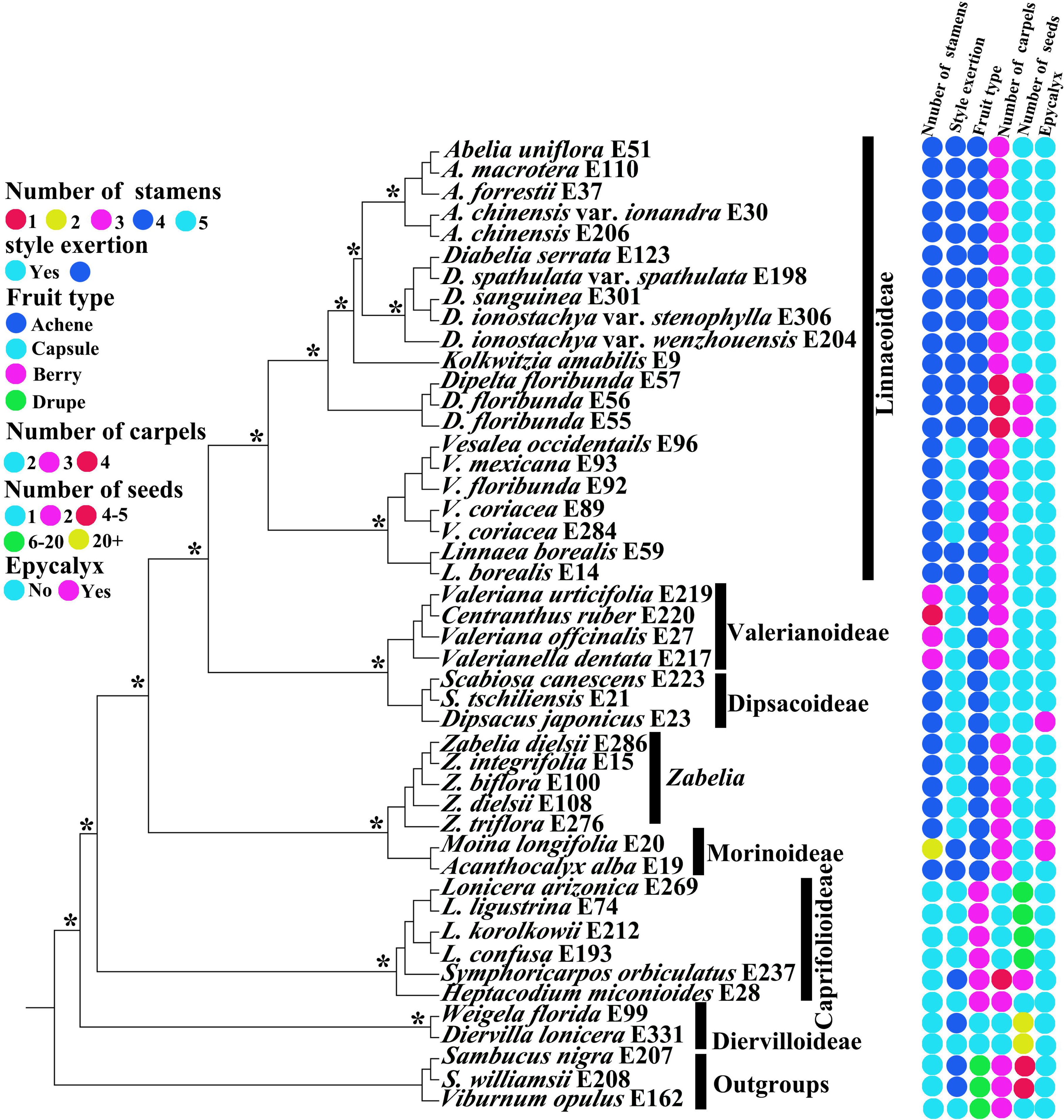
Simplified ML tree generated from the nuclear gene data showing the distribution of selected character states. The asterisks indicate maximum likelihood bootstrap support of 100%.

**Figure. S2.**
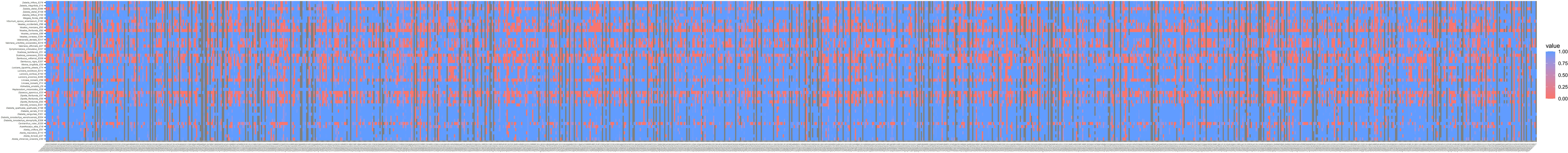
Heatmaps showing gene recovery efficiency for the nuclear gene in 46 species of Caprifoliaceae. Columns represent genes, and each row is one sample. Shading indicates the percentage of the reference locus length coverage.

**Fig. S3.**
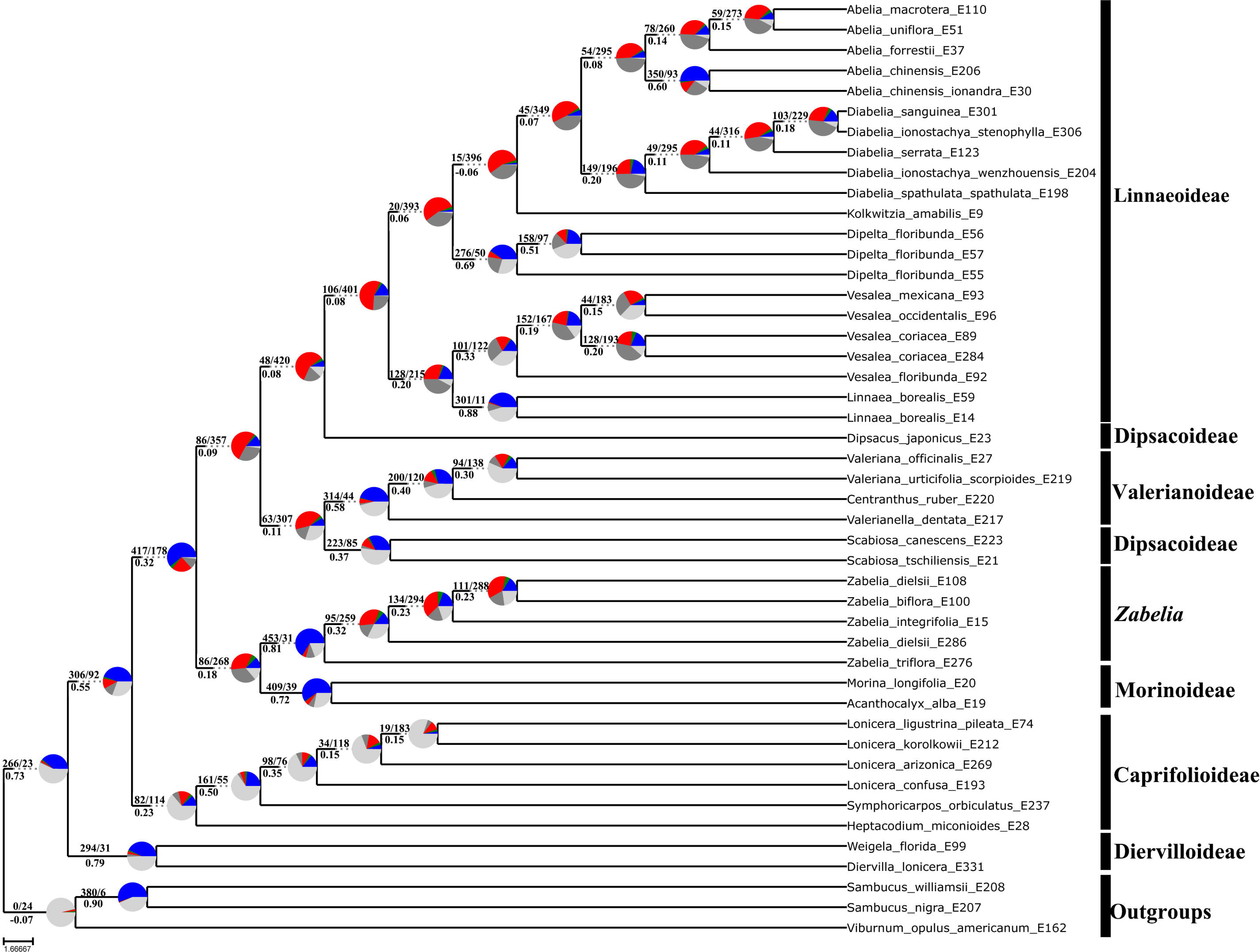
ASTRAL-III species tree. Numbers above branches indicate the number of gene trees concordant/conflicting with that node in the species tree. Numbers below the branches are the Internode Certainty All (ICA) score. Pie charts next to the nodes present the proportion of gene trees that supports that clade (blue), the proportion that supports the main alternative for that clade (green), the proportion that supports the remaining alternatives (red), light gray means missing data, and dark gray mean uninformative (BS < 50%).

**Fig. S4.**
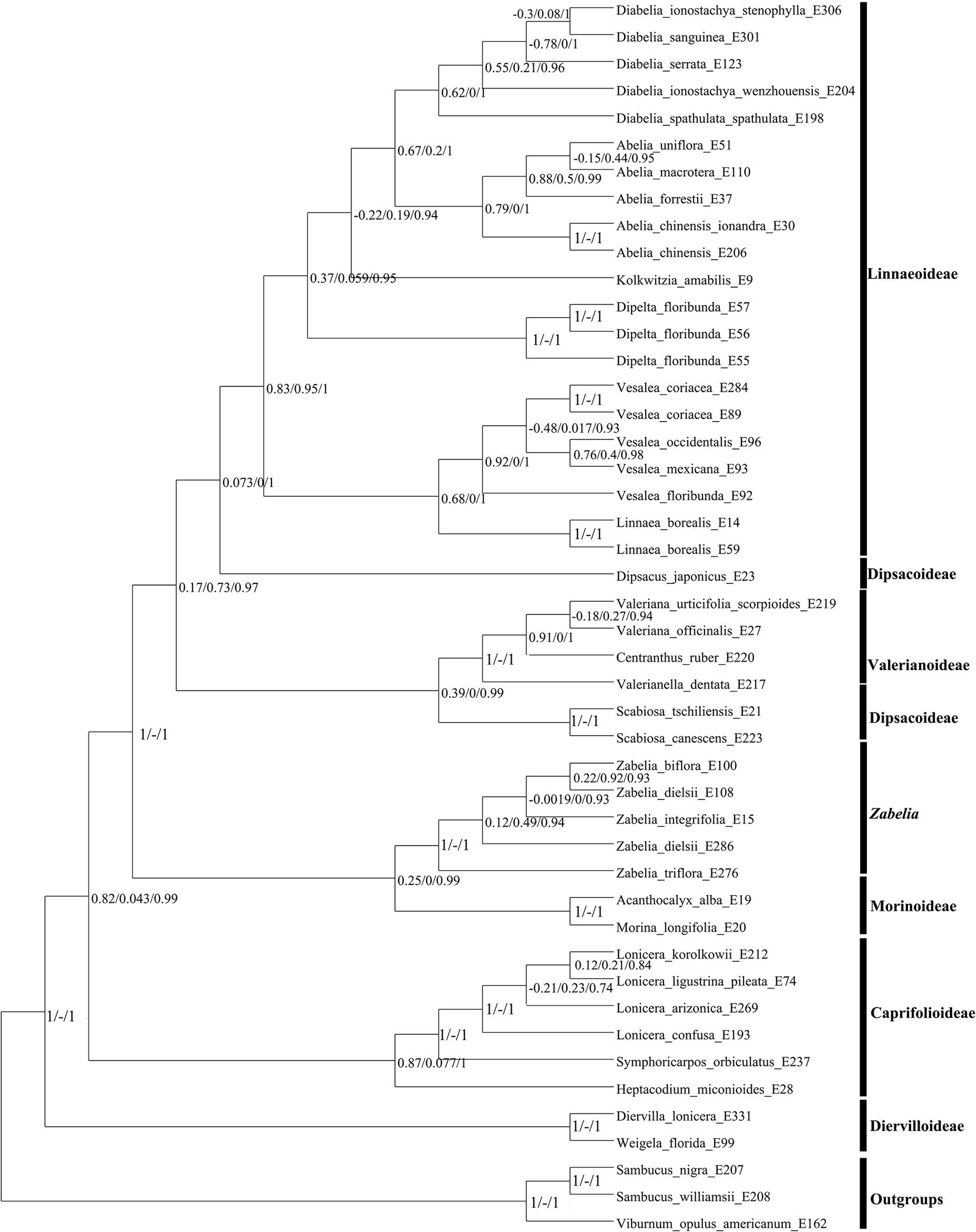
Results of the Quartet Sampling of the ASTRAL tree. Node labels indicate QC/Quartet Differential (QD)/Quarte Informativeness (QI) scores.

**Fig. S5.**
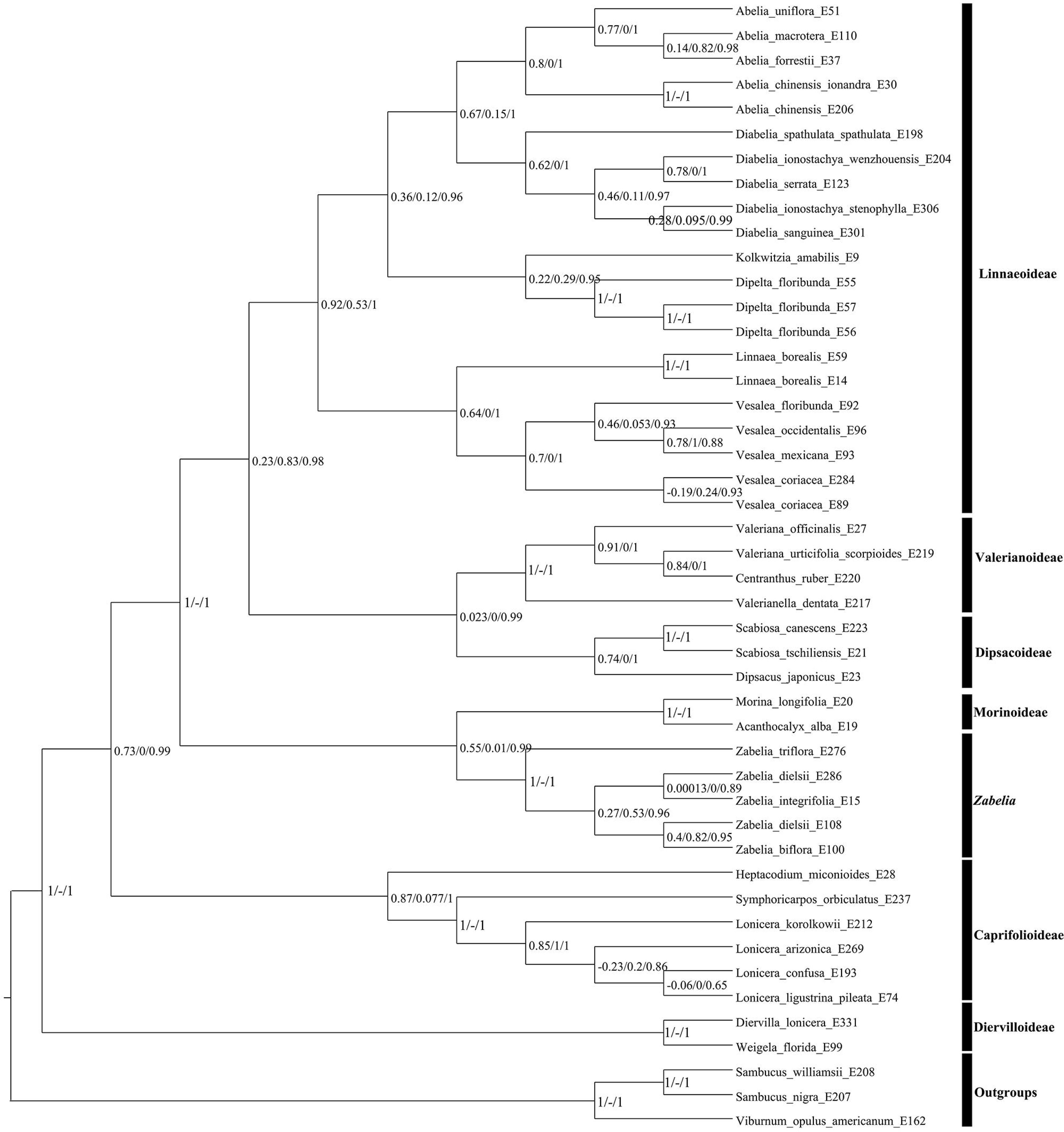
Results of the Quartet Sampling of the nuclear concatenated RAxML tree. Node labels indicate QC/Quartet Differential (QD)/Quarte Informativeness (QI) scores.

**Fig. S6.**
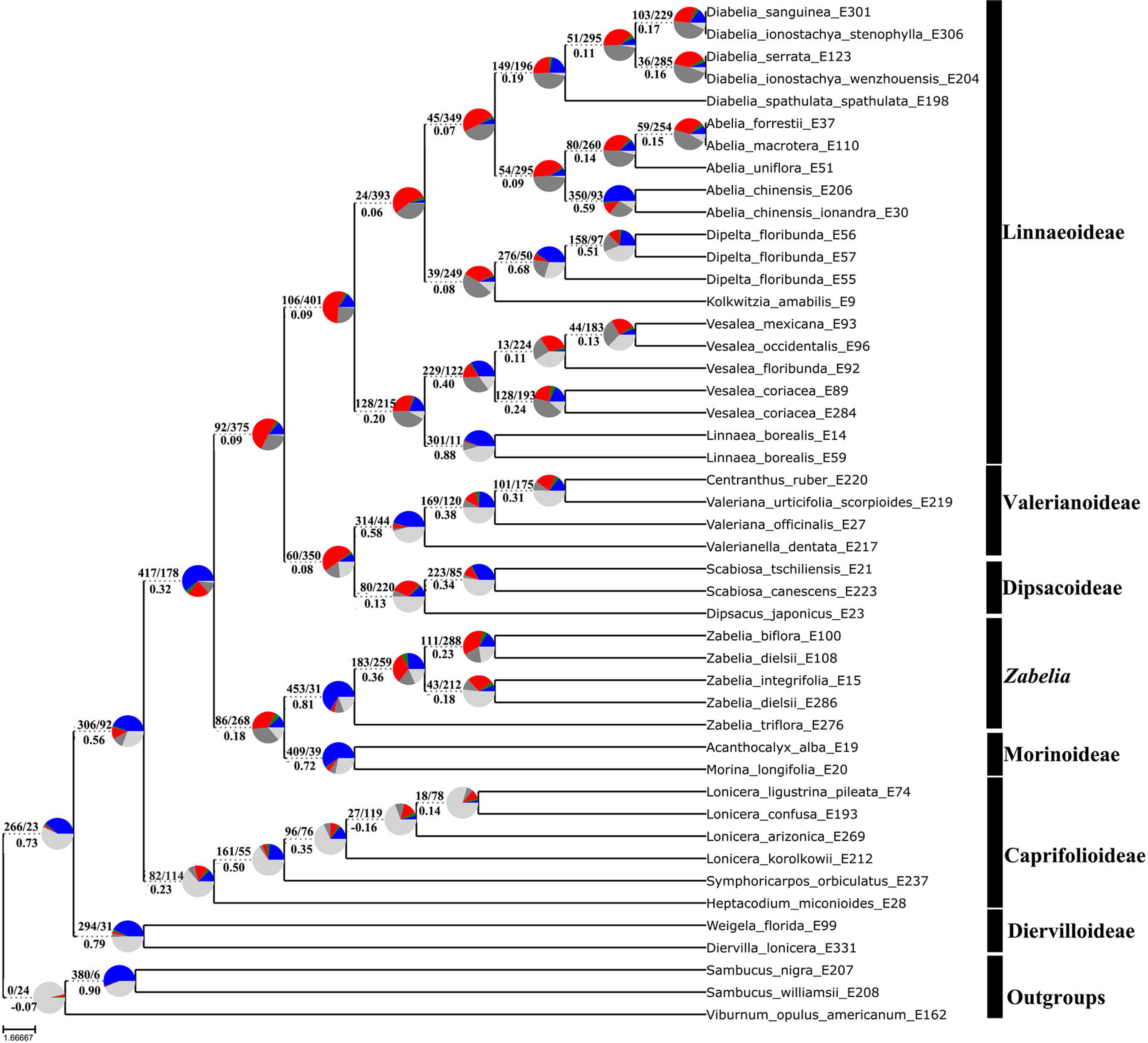
Maximum likelihood cladogram of Caprifoliaceae inferred from RAxML analysis of the concatenated 713-nuclear gene supermatrix. Numbers above branches indicate the number of gene trees concordant/conflicting with that node in the species tree. Numbers below the branches are the Internode Certainty All (ICA) score. Pie charts next to the nodes present the proportion of gene trees that supports that clade (blue), the proportion that supports the main alternative for that clade (green), the proportion that supports the remaining alternatives (red), light gray means missing data, and dark gray mean uninformative (BS < 50%).

**Fig. S7.**
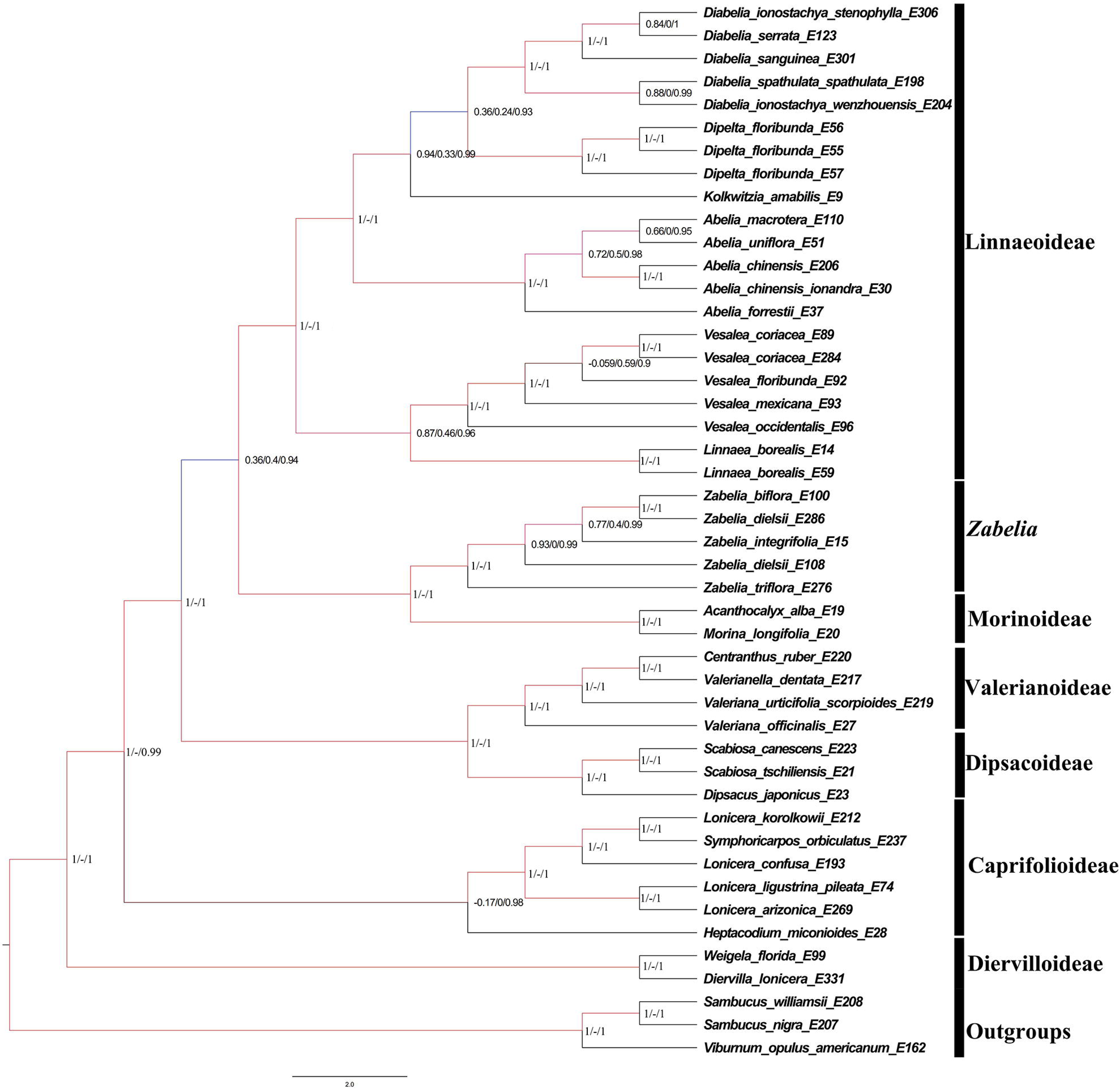
Results of the Quartet Sampling of the chloroplast tree. Node labels indicate QC/Quartet Differential (QD)/Quarte Informativeness (QI) scores.

**Fig. S8.**
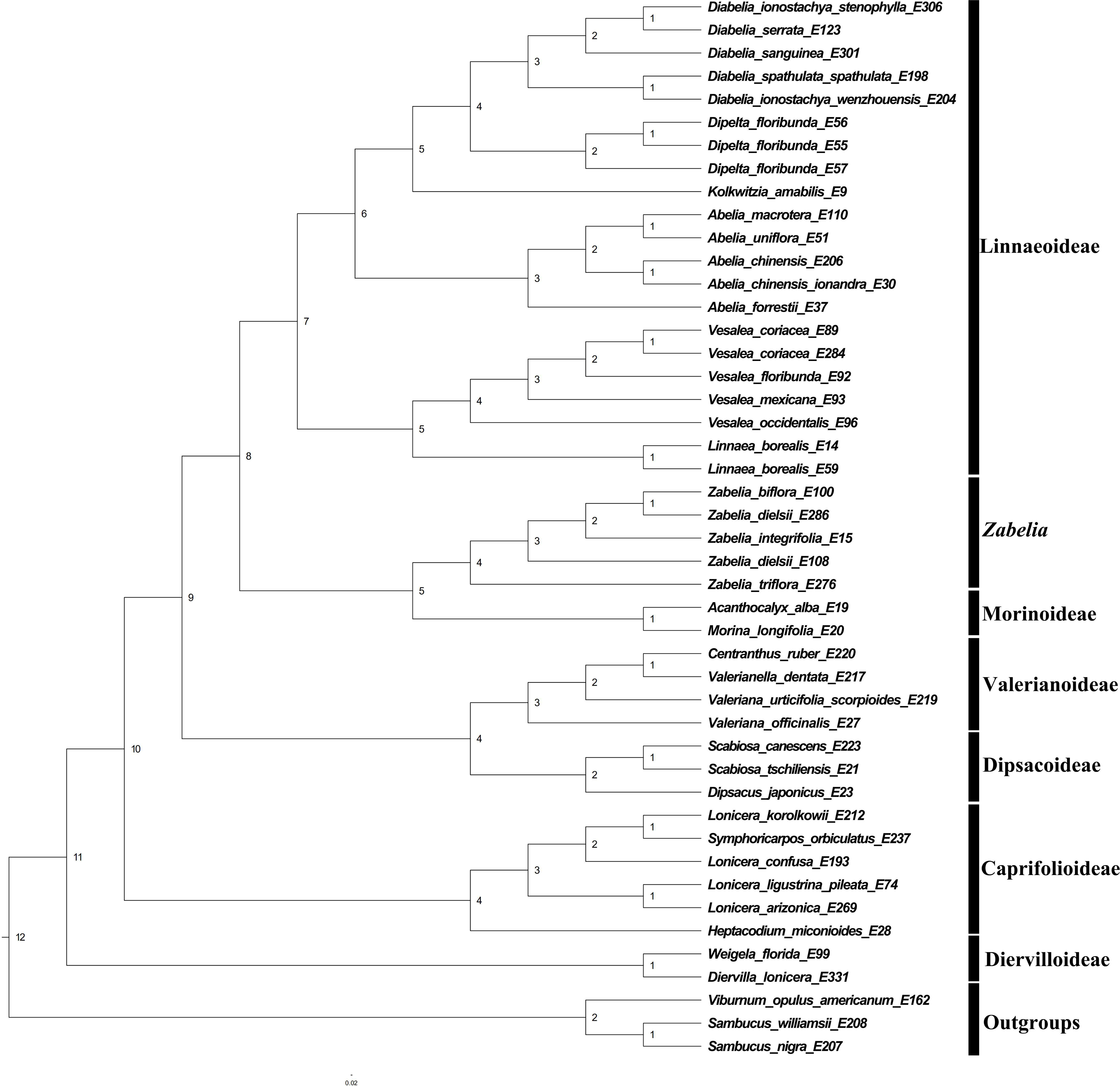
Phylogeny of the plastid DNA dataset; numbers above branches represent clade frequencies of the simulated gene trees.

**Fig. S9.**
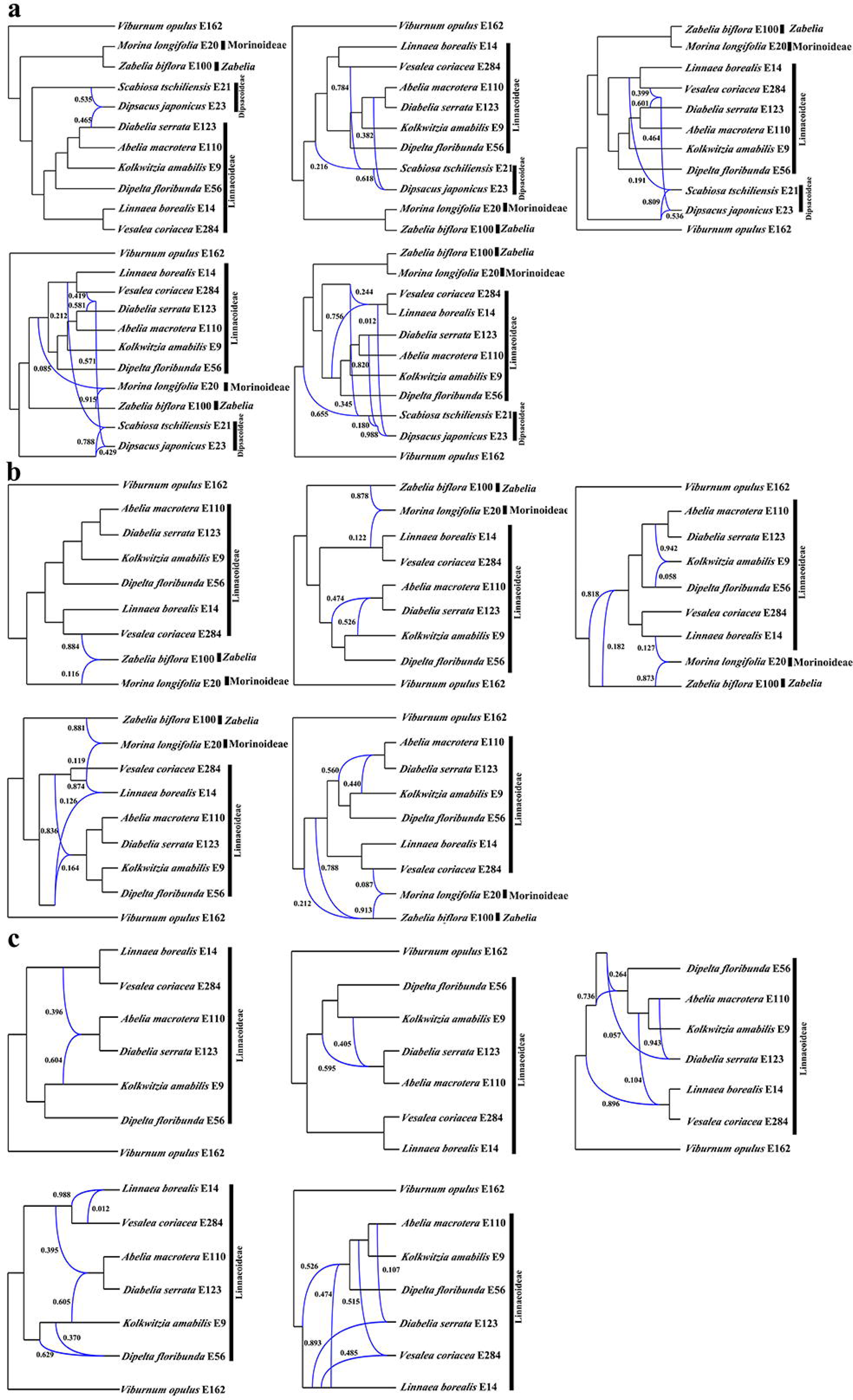
Best supported species networks inferred with Phylonet for the (a) 11-taxa, (b) 9-taxa, and (c) 7-taxa data sets. Blue branches connect the hybrid nodes. Numbers next to the hybrid branches indicate inheritance probabilities.

**Fig. S10.**
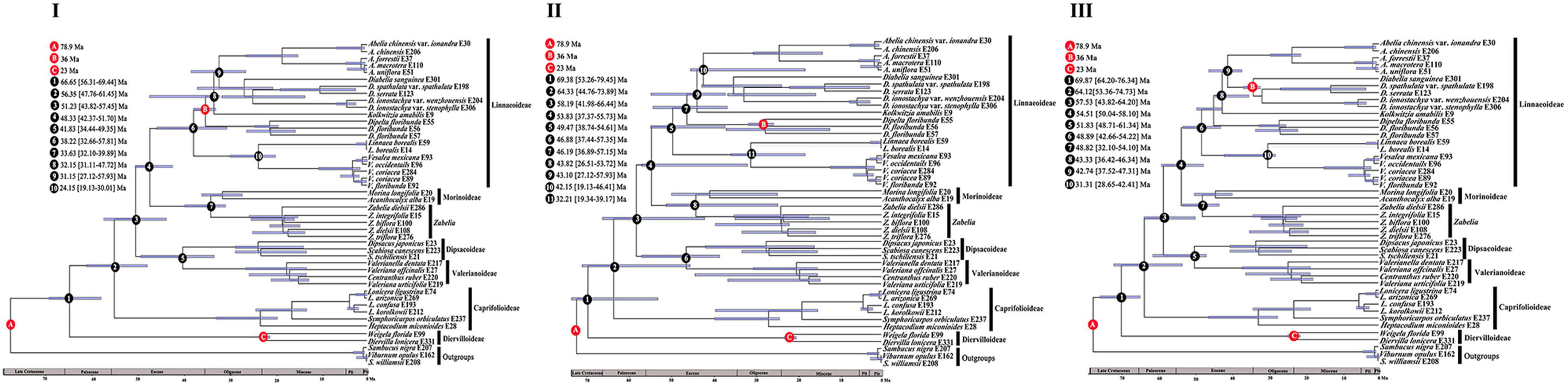
BEAST analysis of divergence times based on the nuclear data. Calibration points are indicated by A, B. and C. Numbers 1–11 represent major divergence events in Caprifoliaceae; mean divergence times and 95% highest posterior densities are provided for each. □, □ and □ indicate the three analyses that varied in the placement of the *Diplodipelta* fossil constraint.

**Fig. S11.**
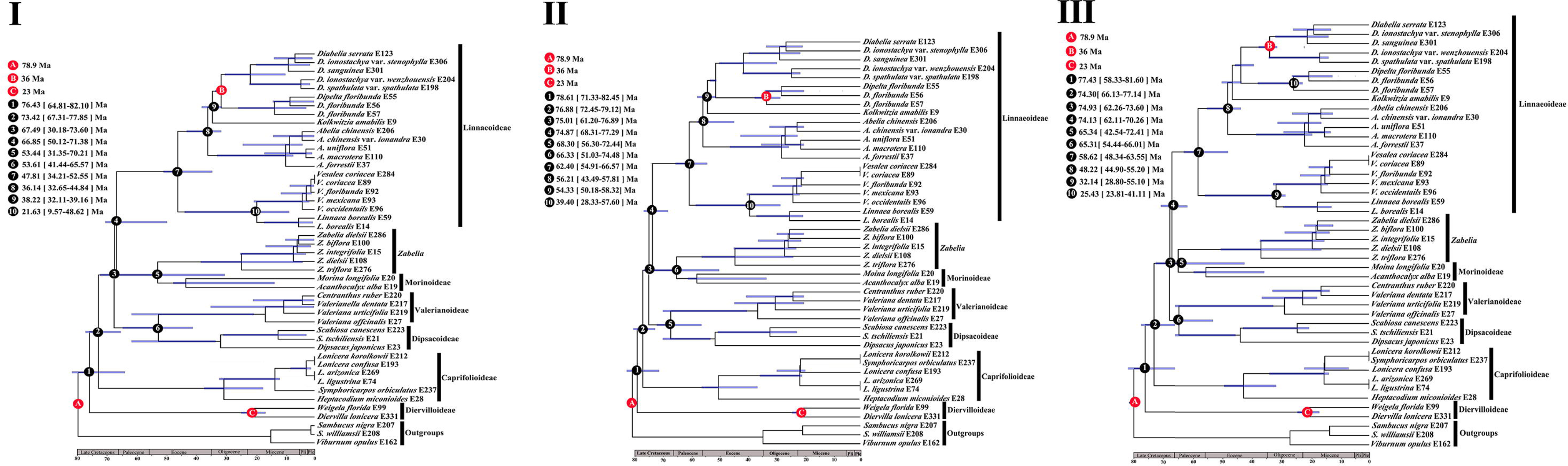
BEAST analysis of divergence times based on the cpDNA data. Calibration points are indicated by A, B, and C. Numbers 1–10 represent major divergence events in Caprifoliaceae; mean divergence times and 95% highest posterior densities are provided for each. □, □ and □ indicate the three analyses that varied in the placement of the *Diplodipelta* fossil constraint.

**Fig. S12.**
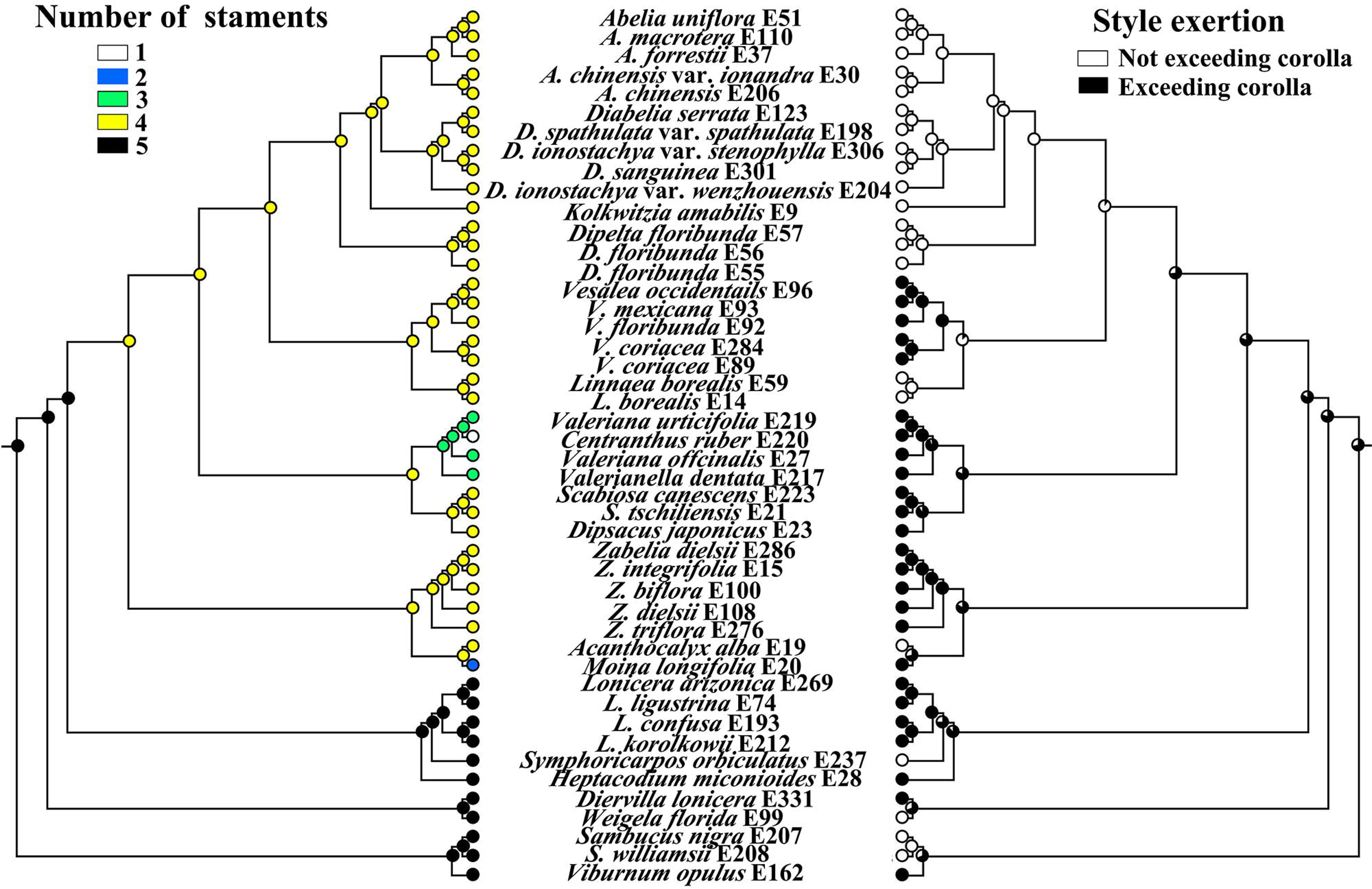
Maximum likelihood inference of character evolution in Caprifoliaceae based on the nuclear matrix. Left, Number of stamens; Right, Style exertion.

**Fig. S13.**
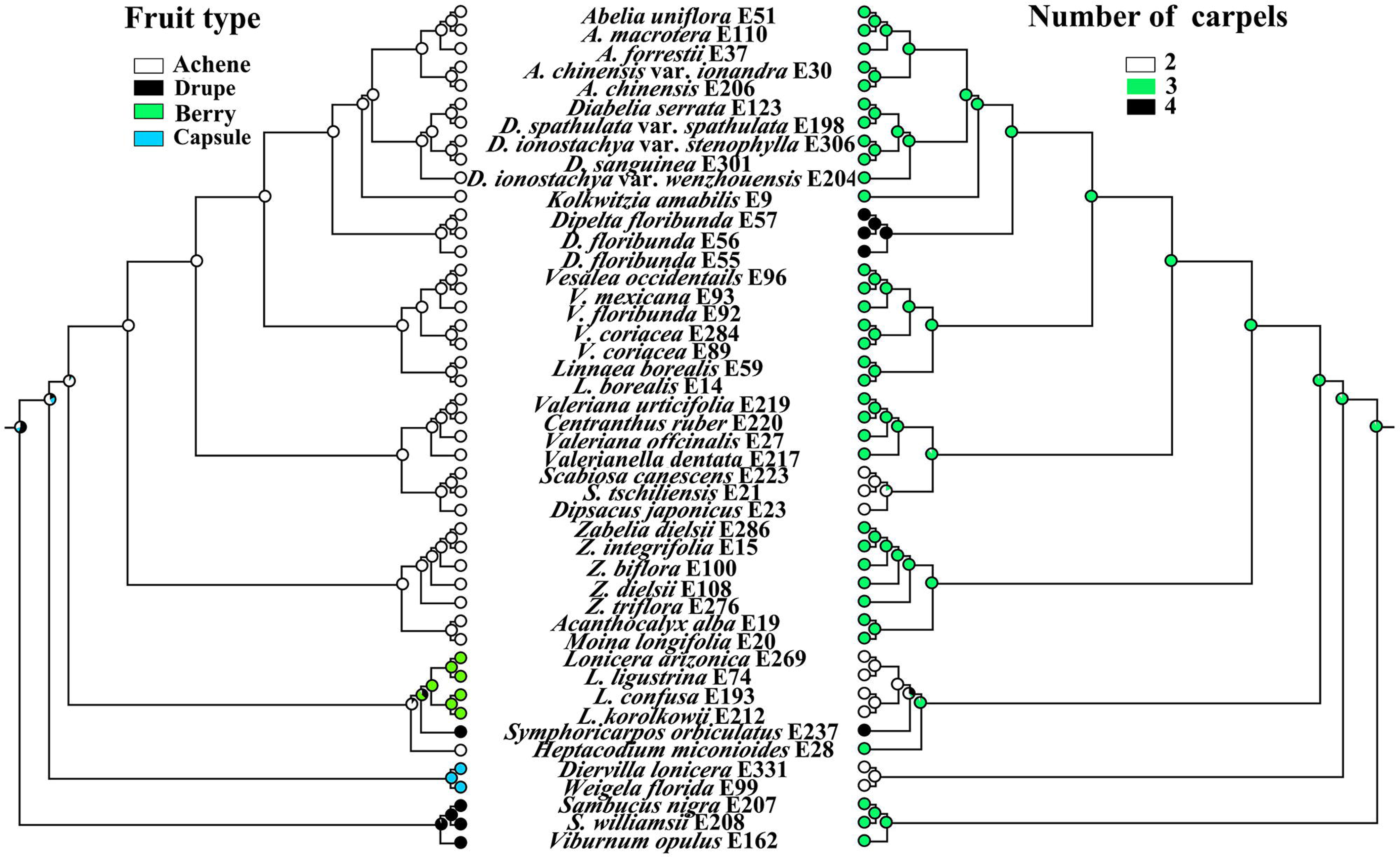
Maximum likelihood inference of character evolution in Caprifoliaceae based on the nuclear matrix. Left, Style of fruit; Right, Number of carpels.

**Fig. S14.**
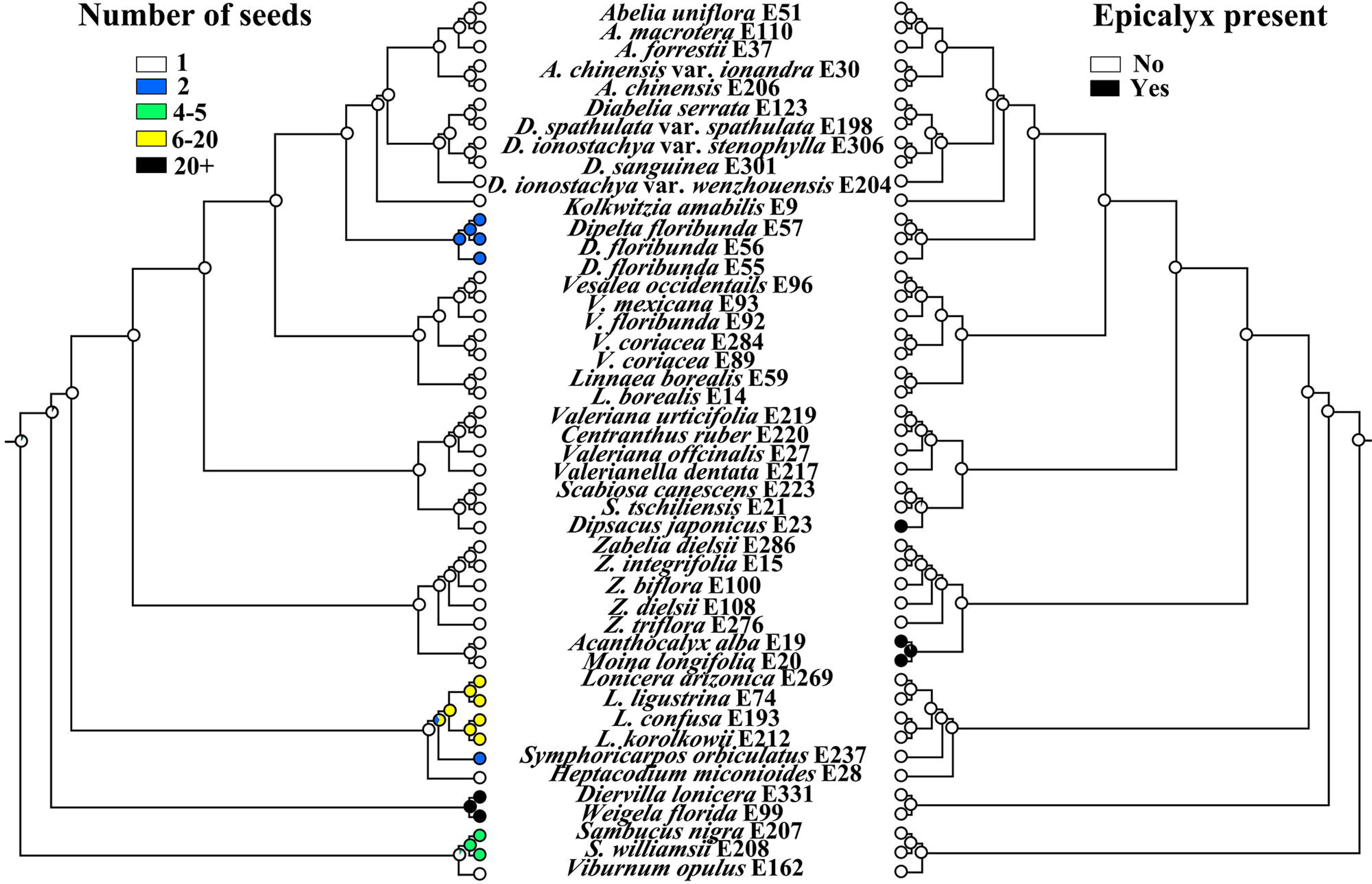
Maximum likelihood inference of character evolution in Caprifoliaceae based on the nuclear matrix. Left, number of seeds; Right, epicalyx presence/absence.

**Table S1.**
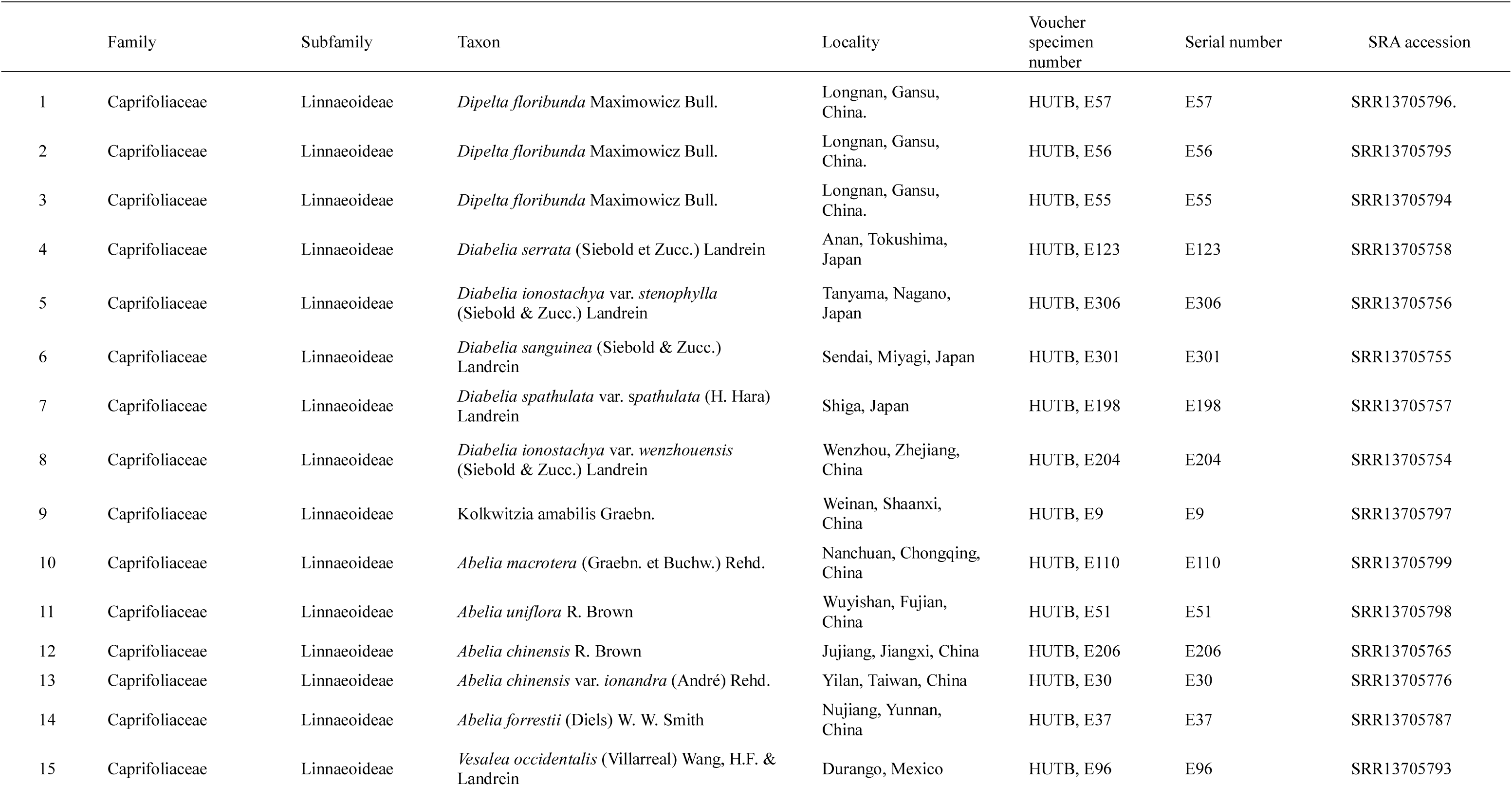
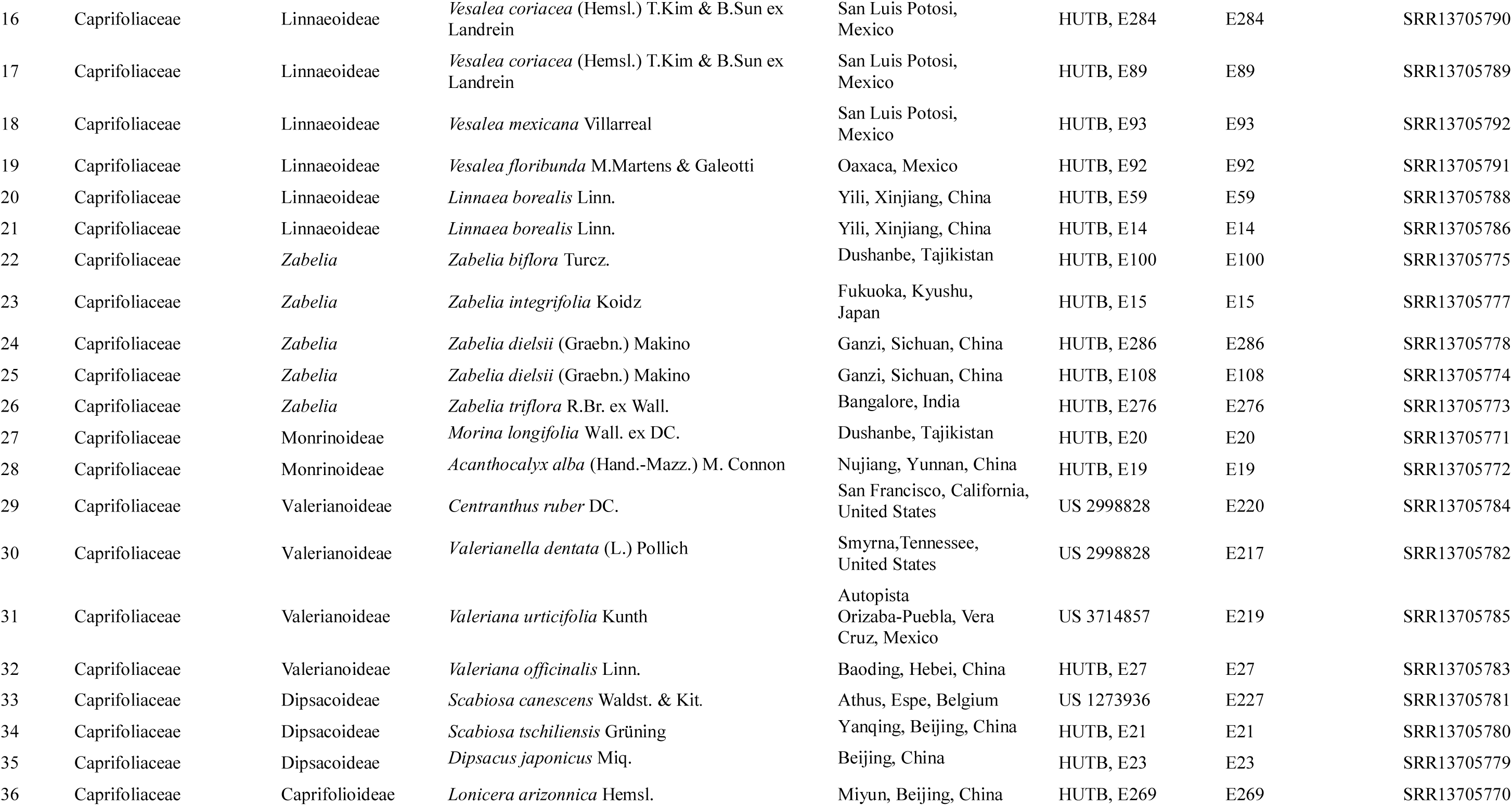
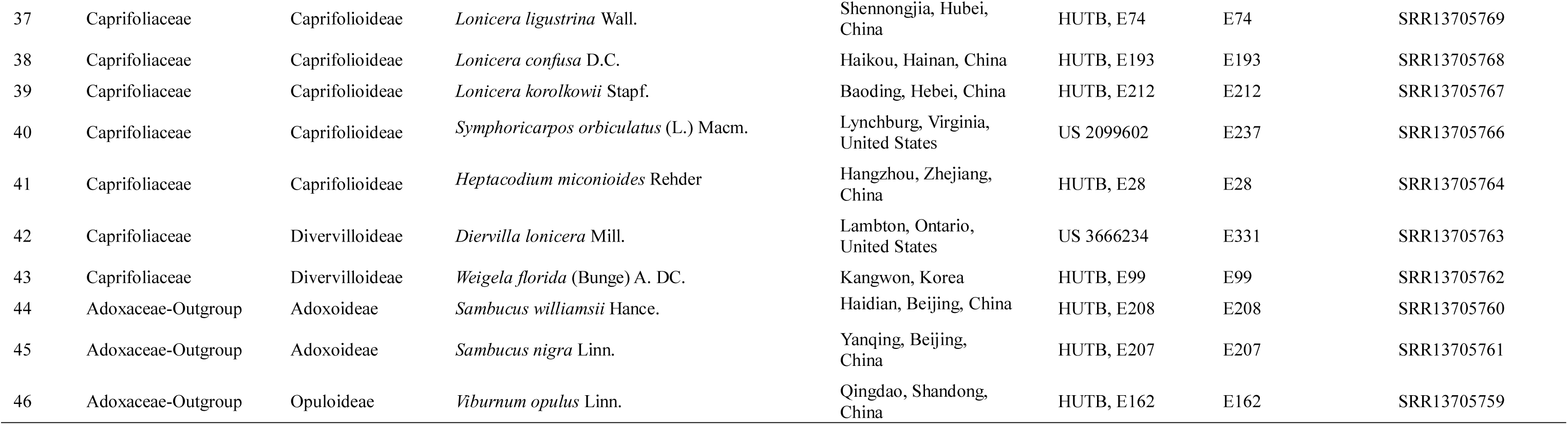
List of species and vouchers used in this study.

**Table S2.**
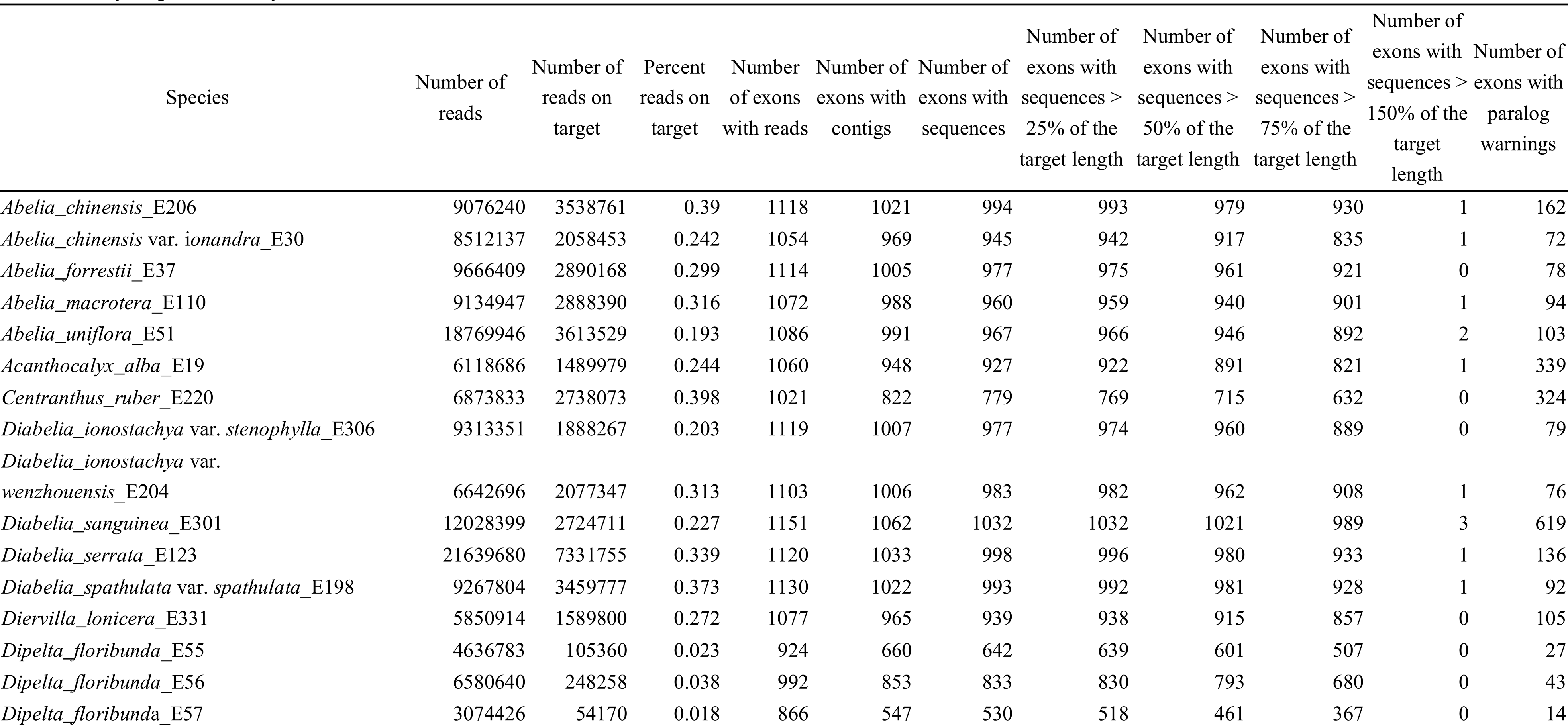
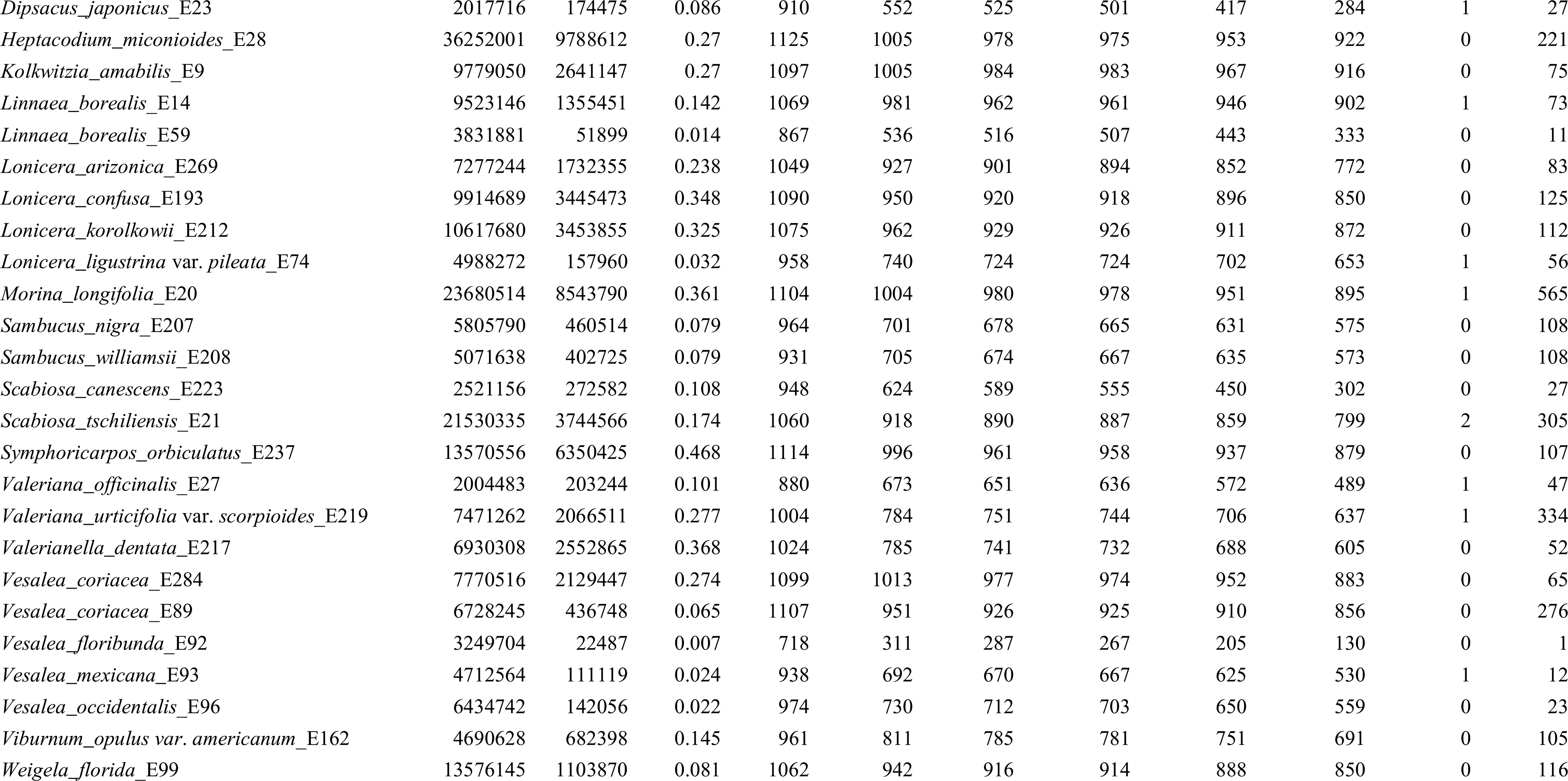
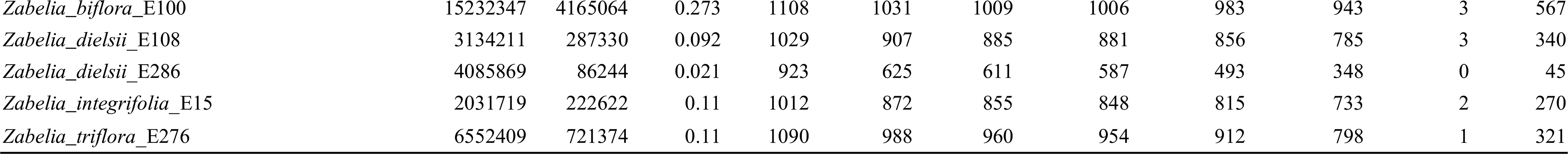
HybPiper assembly statistics

